# Single-cell sequencing reveals clonally expanded plasma cells during chronic viral infection produce virus-specific and cross-reactive antibodies

**DOI:** 10.1101/2021.01.29.428852

**Authors:** Daniel Neumeier, Alessandro Pedrioli, Alessandro Genovese, Ioana Sandu, Roy Ehling, Kai-Lin Hong, Chrysa Papadopoulou, Andreas Agrafiotis, Raphael Kuhn, Damiano Robbiani, Jiami Han, Laura Hauri, Lucia Csepregi, Victor Greiff, Doron Merkler, Sai T. Reddy, Annette Oxenius, Alexander Yermanos

## Abstract

Plasma cells and their secreted antibodies play a central role in the long-term protection against chronic viral infection. However, due to experimental limitations, a comprehensive description of linked genotypic, phenotypic, and antibody repertoire features of plasma cells (gene expression, clonal frequency, virus specificity, and affinity) has been challenging to obtain. To address this, we performed single-cell transcriptome and antibody repertoire sequencing of the murine bone marrow plasma cell population following chronic lymphocytic choriomeningitis virus infection. Our single-cell sequencing approach recovered full-length and paired heavy and light chain sequence information for thousands of plasma cells and enabled us to perform recombinant antibody expression and specificity screening. Antibody repertoire analysis revealed that, relative to protein immunization, chronic infection led to increased levels of clonal expansion, class-switching, and somatic variants. Furthermore, antibodies from the highly expanded and class-switched (IgG) plasma cells were found to be specific for multiple viral antigens and a subset of clones exhibited cross-reactivity to non-viral- and auto-antigens. Integrating single-cell transcriptome data with antibody specificity suggested that plasma cell transcriptional phenotype was correlated to viral antigen specificity. Our findings demonstrate that chronic viral infection can induce and sustain plasma cell clonal expansion, combined with significant somatic hypermutation, and can generate cross-reactive antibodies.

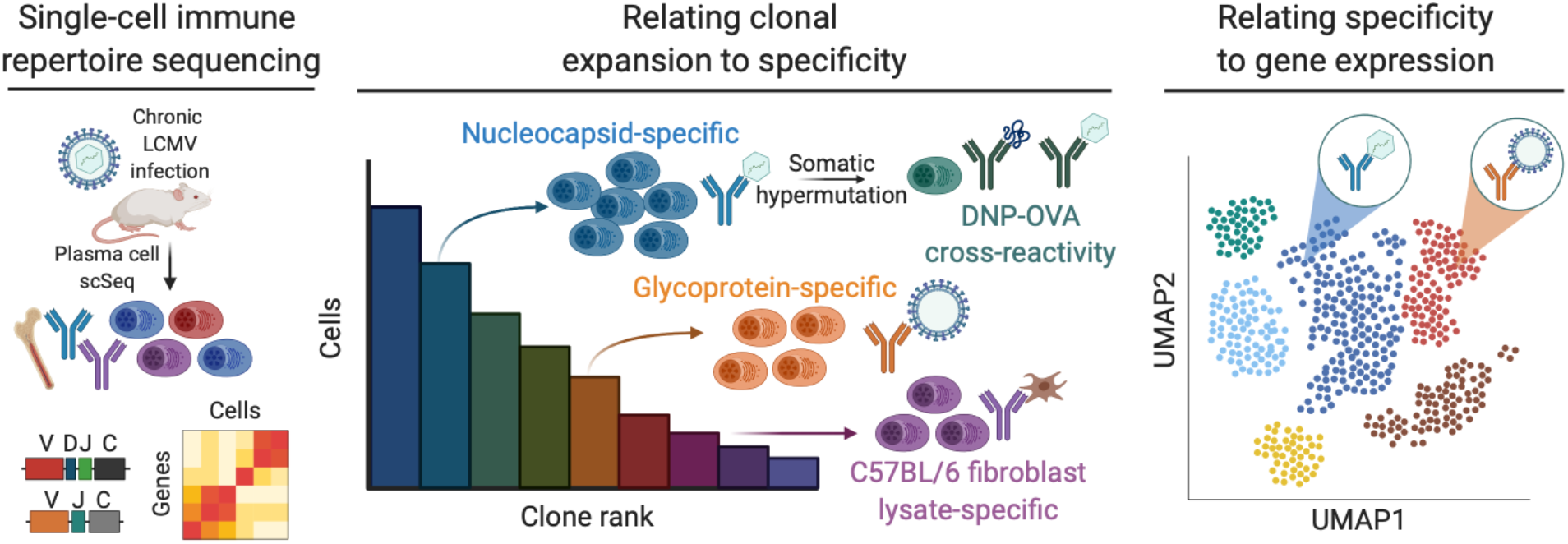
Graphical abstract. Single-cell sequencing reveals clonally expanded plasma cells during chronic viral infection produce virus-specific and cross-reactive antibodies.

## Introduction

The humoral immune response plays a critical role in defending the host from a variety of pathogens, whereby the terminally differentiated plasma cell population contributes by secreting antibodies that can clear current and prevent future infection (Bootz et al., 2017; Burton and Hangartner, 2016). Lymphocytic choriomeningitis virus (LCMV) represents a well-studied pathogen that can establish chronic (dose-dependent) infections in mice. Such infections can eventually be controlled and cleared due to the emergence of plasma cells secreting neutralizing antibodies. Circulating virus-neutralizing antibodies are detected in the serum, but their plasma cell producers are mostly restricted to lymphoid tissues (Greczmiel et al., 2017; Nguyen et al., 2019). It has further been established that neutralizing antibodies of the class-switched IgG isotype are especially crucial for the resolution of chronic LCMV infection (Battegay et al., 1993; Hangartner et al., 2003). These neutralizing IgG antibodies are directed against the LCMV surface glycoprotein complex (GPC) and only emerge several months after initial infection. In addition to neutralizing GPC-targeting antibodies, GPC-specific non-neutralizing antibodies and antibodies targeting the nucleocapsid protein (NP) are also observed shortly after infection (Battegay et al., 1993; Kimmig and Lehmann-Grube, 1979). Moreover, in addition to LCMV-specific antibodies, it has been shown that chronic LCMV infection in mice also induces polyclonal hypergammaglobulinemia (PHGG), which induces production of LCMV-unspecific IgG antibodies. Induction of PHGG is mediated by LCMV-specific non-follicular CD4 helper cells, which support the activation of B cells in a B-cell receptor independent manner. For instance, LCMV infection results in plasma cells producing antibodies specific to unrelated antigens such as ovalbumin and dinitrophenol (DNP) or autoantigens (i.e., double- and single-stranded DNA, insulin or thyroglobulin) (Greczmiel et al., 2020; Hunziker et al., 2003), whereby these autoantibody titers can even be comparable to the concomitantly induced antiviral antibody response (Ludewig et al., 2004).

The bone marrow represents a stable niche for long-lived plasma cells (BM PCs) (Manz et al., 1997); in this terminal differentiation state, BM PCs have lost surface antibody (B cell receptor) expression in exchange for an increased antibody secretion capacity (Halliley et al., 2015; Kräutler et al., 2020; Medina et al., 2002). This lack of antibody surface expression complicates the identification and subsequent isolation of virus-specific and class-switched plasma cells by means of traditional fluorescence-activated cell sorting (FACS) methods. It has previously been shown that LCMV-specific plasma cells reside in the bone marrow following infection (Kräutler et al., 2020) via ELISpot assays (Painter et al., 2014; Wolf et al., 2011), in which antigen-specificity for a given target can be quantified. However, such assays do not allow for recovery of the underlying antibody sequences and therefore information regarding the relationship between clonal expansion, somatic hypermutation, and affinity to viral antigens cannot be obtained. Furthermore, these assays do not allow a detailed transcriptional characterization of individual antibody-secreting cells.

Recent advances in single-cell sequencing (scSeq) technologies have made it possible to obtain both transcriptome and antibody repertoire information from single B cells at large scale (Croote et al., 2018; Horns et al., 2020; Saikia et al., 2019; Singh et al., 2019) enabling an integrated analysis of genotypic and phenotypic metrics of paired antibody repertoires.

Here, we performed scSeq to molecularly quantify both the antibody repertoire and transcriptome of thousands of murine BM PCs from a mouse following chronic viral infection and compare this with mice immunized with protein antigens. Importantly, we also took advantage of scSeq data to reconstruct full-length antibody proteins from BM PCs in order to screen for antigen specificity. Such extensive scSeq combined with experimental antibody characterization (synthetic gene construction, cloning, recombinant expression and antigen-binding assays) limited our capacity to perform this on only a single chronically infected LCMV mouse. Nevertheless, the single-cell linking of molecular genotype, phenotype, and antigen specificity of B cells has rarely been achieved before. One of the only examples to date has been in the context of human influenza vaccination and was also confined to a single individual (Horns et al., 2020). We report that relative to protein immunization, chronic viral infection resulted in an increased proportion of clonally expanded, class-switched, and somatically hypermutated BM PCs. We could additionally demonstrate that highly expanded IgG-secreting bone marrow plasma cells produce virus-specific, cross-reactive, and potentially autoreactive antibodies. Finally, single-cell transcriptome sequencing analysis suggests that plasma cells producing antibodies specific for the LCMV GPC protein occupy distinct transcriptional states relative to plasma cells secreting NP-specific antibodies. Together, our findings inform the relationship between clonal expansion, somatic hypermutation, gene expression and antigen-specificity during chronic LCMV infection.

## Results

### Chronic viral infection results in high clonal expansion of plasma cells expressing class-switched IgG

To molecularly characterize the antibody repertoire and gene expression profiles following chronic viral infection, we isolated BM PCs 28 days post infection (dpi) with high-dose (2×10^6^ focus forming units (ffu)) LCMV clone 13 (Figure 1A). We performed FACS to isolate BM PCs based on CD138+, TACI+ CD19^lo^, and B220^lo^ expression (as previously described) (Pracht et al., 2017) and performed scSeq of transcriptomes and antibody repertoires using the 5’ GEX and V(D)J protocols from 10X Genomics. Additionally, we performed scSeq of BM PC from mice that had undergone serial immunizations with the protein antigens ovalbumin (OVA) or the extracellular domain of human tumor necrosis factor receptor 2 (TNFR2; 61% amino acid sequence homology to its murine homolog); this allowed us to compare transcriptome and repertoire features of chronic infection to protein immunization (Figures S1A, S1B). We were able to recover thousands of cells for each mouse (Figures S1C, S1D), with chronic LCMV infection resulting in the highest proportion of class-switched IgG producing BM PCs and clones (Figures 1B, 1C, S1E, S1F): we detected 5337 IgG-expressing BM PCs (Figure 1B) corresponding to 1653 unique clones (Figure 1C). Unique clones are defined herein as cells expressing only one heavy and one light chain and containing identical complementarity determining region 3 variable heavy and complementarity determining region 3 variable light chain (CDRH3+CDRL3) amino acid sequences. The discrepancy between the number of cells and the number of unique clones for all mice and isotypes suggested significant clonal expansion in the BM PC repertoires. Upon closer inspection of the 50 most expanded clones in each mouse (Figure 1D), we observed a striking difference in the proportion of IgG expressing BM PCs: 46 clones were expressing IgG, with 10 clones showing high clonal expansion (composed of more than 50 distinct cells) in chronic virus-infected mice compared to only four IgG expressing clones, with less clonal expansion in both protein immunized mice (Figure 1D).

**Figure 1.**
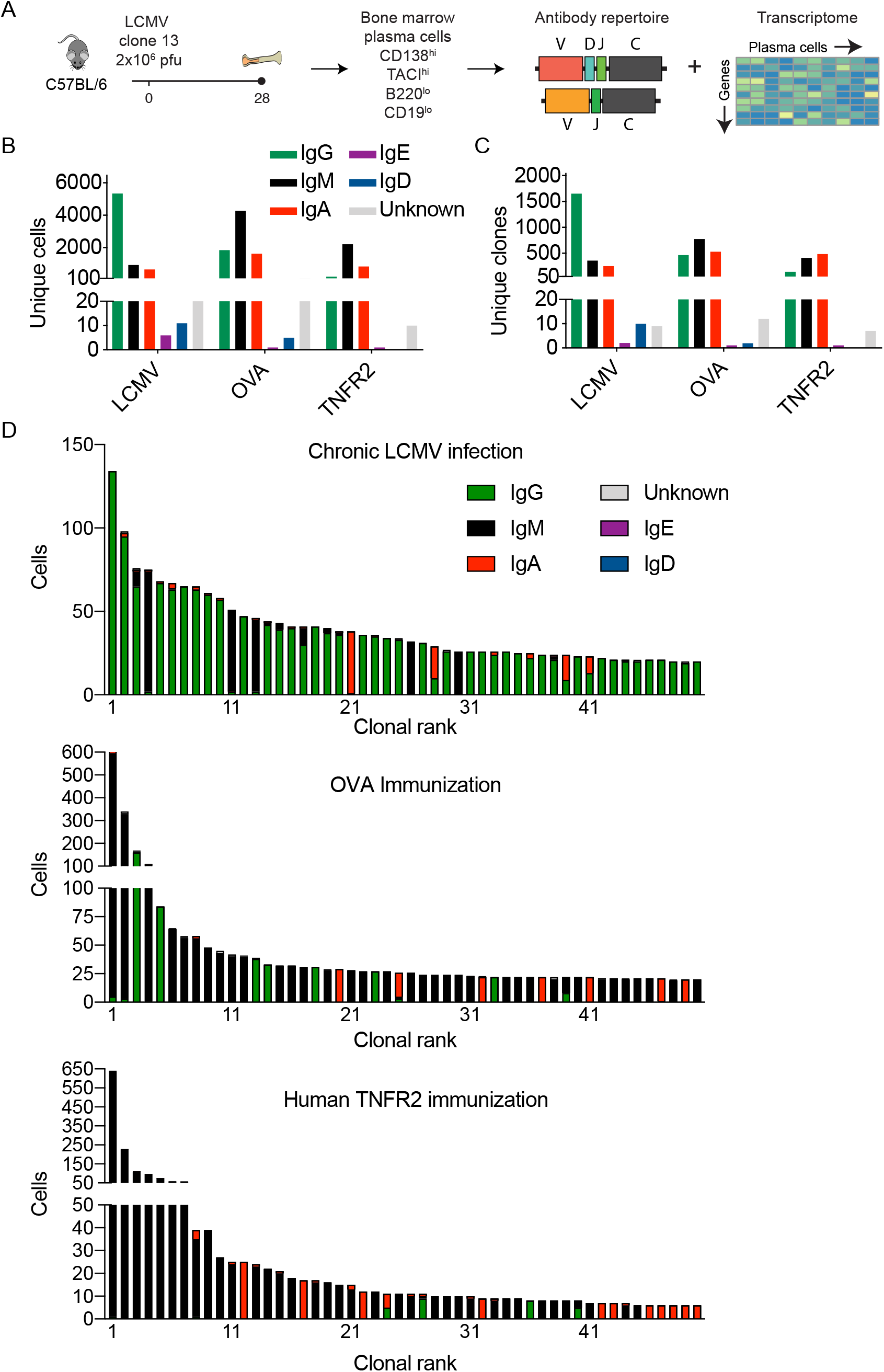
Single-cell immune repertoire sequencing reveals clonal expansion and class-switching in the bone marrow plasma cell repertoire following chronic viral infection. A. Experimental overview of chronic LCMV infection, bone marrow plasma cell isolation, and single-cell sequencing. B. Number of cells per isotype for each infected or immunized mouse. Only cells containing exactly one variable heavy (V_H_) and variable light (V_L_) chain were considered. Colors correspond to isotype. C. Number of clones per isotype for each infected or immunized mouse. Clones were determined by grouping those B cells containing identical CDRH3+CDRL3 amino acid sequences. The isotype was determined as the isotype corresponding to the majority of cells within one clone. Color corresponds to isotype. D. Clonal expansion for the top 50 most expanded clones of the bone marrow plasma cells for each infected or immunized mouse. Clones were determined by grouping those B cells containing identical CDRH3+CDRL3 amino acid sequences. Color corresponds to isotype.

Given that the majority of clonally expanded BM PCs expressed IgG during chronic LCMV infection, we next determined the fraction that expressed the IgG2c subtype, as IgG2c is the most dominant isotype of LCMV-specific antibodies and has previously been shown to be crucial for viral neutralization (Barnett et al., 2016). Further analysis of the expanded IgG clones revealed heterogeneous expression of IgG1, IgG2b, and IgG2c following chronic viral infection, but not following protein immunizations (Figures 2A, S2A, S2B). Interestingly, in the case of chronic viral infection, clones predominantly expressing IgG1 did not contain many cells producing the other IgG subtypes, whereas clones predominantly expressing IgG2b and IgG2c contained more variation in subtypes. Quantifying the clonal expansion across the three BM PC repertoires suggested that chronic viral infection increased both the proportion and number of clonally expanded cells expressing IgG, coinciding with a decrease in the proportion and number of clonally expanded IgM cells relative to protein immunizations (Figures 2B, 2C).

**Figure 2.**
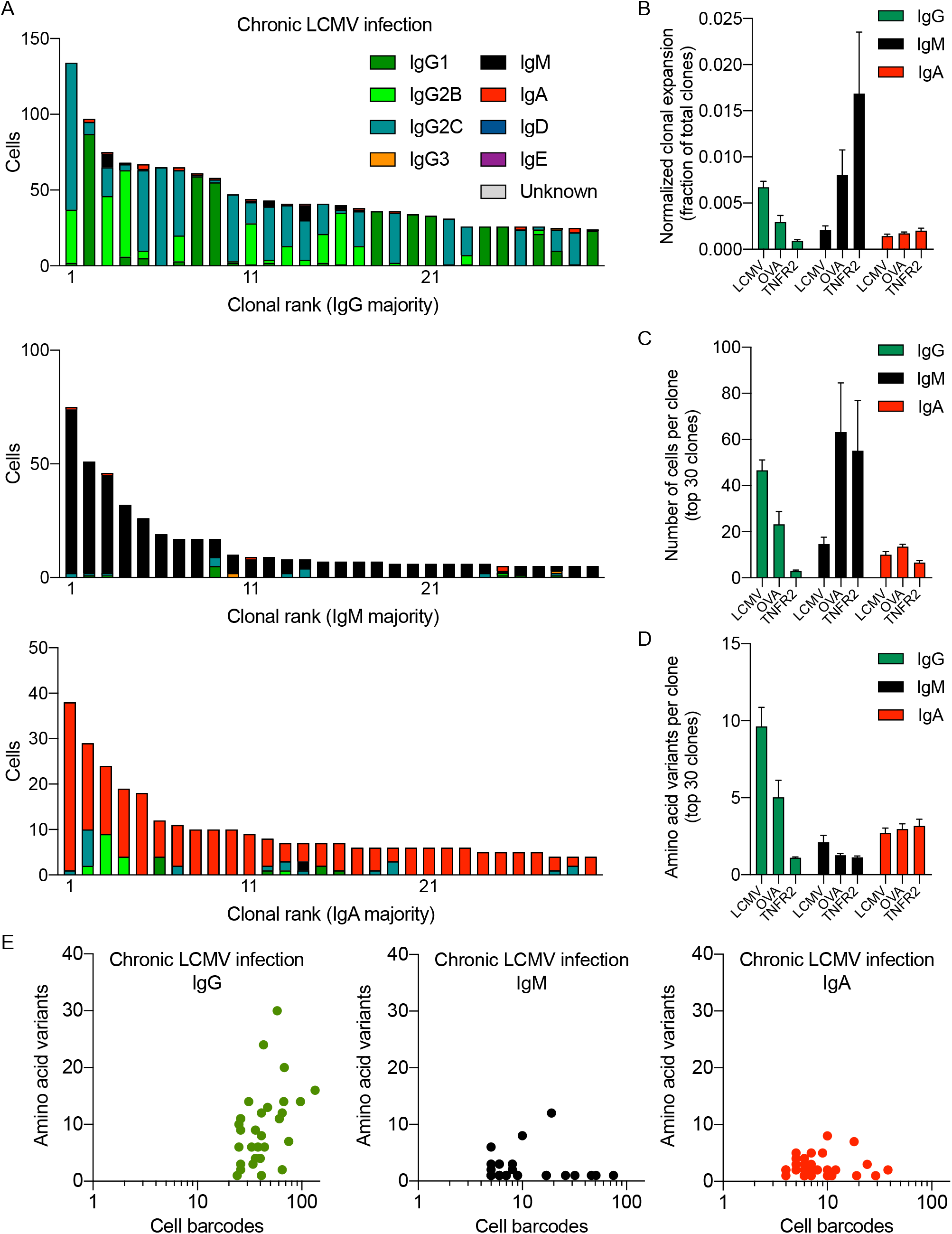
Clonal expansion and number of antibody variants in the IgG bone marrow plasma cell repertoire following chronic viral infection as compared to protein immunizations. A. Clonal expansion of the bone marrow (BM) plasma cells (PC) separated by isotype for a single mouse following chronic LCMV infection. The number of distinct cell barcodes belonging to the top 30 clones are shown. Clone was determined by grouping those B cells containing identical CDRH3+CDRL3 amino acid sequences. Only cells containing exactly one variable heavy (V_H_) and variable light (V_L_) chain amino acid sequence were considered. The isotype was determined as the isotype corresponding to the majority of cells within one clone. Color corresponds to isotype. B. Normalized clonal expansion separated by isotype for each of the infected or immunized mice. The number of cell barcodes per clone was divided by the total number of cells in each repertoire. C. Number of cells per clone for the top 30 clones separated by isotype for each of the infected or immunized mice. D. Quantity of amino acid variants within each clone separated by isotype for each of the infected or immunized mice. Amino acid variants were determined by quantifying the number of unique full-length V_H_+V_L_ amino acid sequence variants within each clone. Clone was determined by grouping those B cells containing identical CDRH3+CDRL3 amino acid sequences. E. Relationship between the number of unique amino acid variants and the number of cell barcodes for the top 30 clones separated by isotype majority following chronic LCMV infection.

### Chronic viral infection drives high levels of somatic hypermutation in clonally expanded plasma cells

After observing that the majority of recovered plasma cells were of the IgG isotype, we hypothesized the most expanded IgG clonal families would contain more somatic hypermutation variants than highly expanded IgM and IgA clones following both viral infection and immunization. We quantified the number of somatic variants (defined by unique, full-length amino acid sequence of paired V_H_ and V_L_) for the 30 most expanded clones of each isotype. Our analysis revealed an increased number of somatic variants of the IgG isotype compared to clones of the IgA or IgM isotype for both LCMV infection and OVA immunization, but not for immunization with TNFR2 (Figures 2D, S3). Relating the number of cells to the number of unique antibody variants demonstrated an increased Pearson correlation (r=0.39) for the IgG isotype relative to IgM (r=0.09) and IgA (r=0.04), but nevertheless indicated that the most expanded clones are not necessarily those containing the largest sequence variation (Figure 2E).

### Chronic viral infection results in plasma cells with inflammatory and isotype-specific gene expression signatures

We next leveraged the ability to simultaneously perform whole transcriptome-as well as targeted V_H_/V_L_-repertoire scSeq to determine unique gene expression profiles of PC clones from chronic LCMV infection or TNFR2 immunization (scSeq of transcriptomes for OVA immunized mouse was not performed). We first filtered out all cells not found in both the scSeq antibody repertoire and transcriptome datasets for the virus-infected and immunized mice, which resulted in 6446 and 3069 cells, respectively (Figure 3A). After normalizing and scaling gene expression counts for the two sequencing libraries, we performed unsupervised clustering to group cells with similar transcriptional profiles in an unbiased manner and visualized each cell on two dimensions using uniform manifold approximation projection (UMAP), which resulted in 14 clusters with distinct transcriptional signatures (Figures 3A, S4). Quantifying the cluster membership for each sample confirmed the visual observation from the UMAP that the BM PCs from viral infection occupy almost entirely different clusters than the BM PCs following protein immunization, with minor overlap occurring only in clusters 5 and 9 (Figure 3B). We next computed the differentially expressed genes between the samples, which revealed an upregulation in interferon-related genes (*Ifi27l2a* and *Ifitm3*) and a downregulation of genes involved in Fos and Jun signaling pathways (*Fos*, *Fosb*, *Jun*, *Junb*, *Ubc*) in BM PCs from chronic LCMV infection (Figure 3C). While it is possible that batch effects may play a role in these differences, expression of traditional plasma-cell markers (*Sdc1* encoding CD138, *Tnfrsf13b* encoding TACI, *Slamf7*, *Prdm1* encoding *BLIMP1*) were nevertheless comparable between the two samples (Figure S5A, S5B).

**Figure 3.**
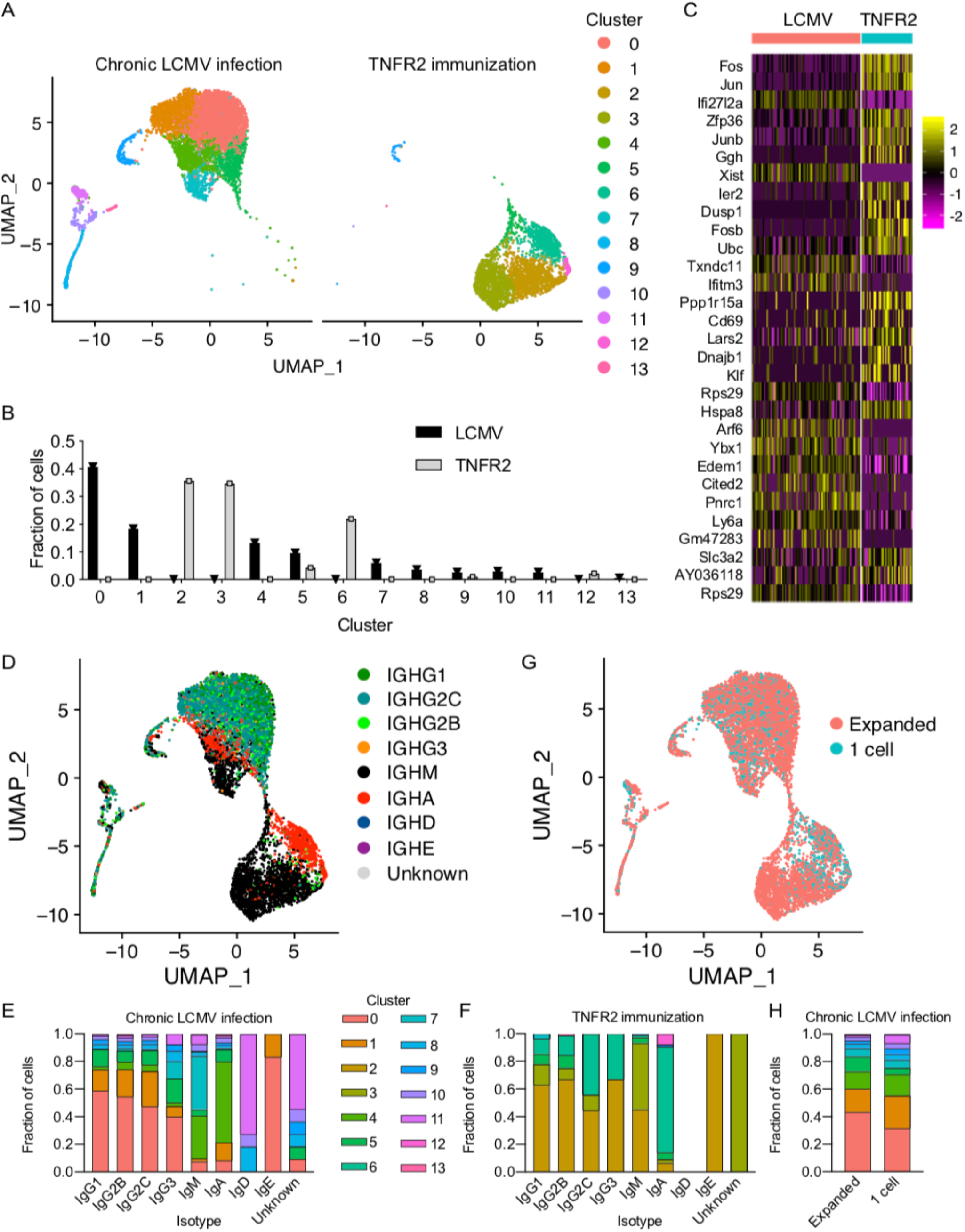
Transcriptional heterogeneity of BM PCs from LCMV-infected or TNFR2-immunized mice. A. Uniform manifold approximation projection (UMAP) based total gene expression for bone marrow plasma cell (BM PC) repertoire following either LCMV infection or TNFR2 immunization. Each point corresponds to a cell and color corresponds to the transcriptional cluster. B. The fraction of cells belonging to each transcriptional cluster from the BM PC repertoire following either LCMV infection or TNFR2 immunization. C. Differentially expressed genes between BM PCs following LCMV infection and TNFR2 immunization. The order of genes (from top to bottom) corresponds to the highest average log fold change. All genes displayed have adjusted p values < 0.01. D. UMAP displaying isotype distribution of BM PCs following either LCMV infection or TNFR2 immunization. Each point corresponds to a cell and color indicates isotype based on VDJ sequencing data. E. Cluster membership separated by isotype for the BM PCs following LCMV infection. Color indicates the transcriptional cluster from panel A. F. Cluster membership separated by isotype for the BM PCs following TNFR2 immunization. Color indicates the transcriptional cluster from panel A. G. UMAP displaying clonal expansion for the BM PCs following either LCMV infection or TNFR2 immunization. “Expanded” corresponds to those clones supported by two or more unique cell barcodes. Clone was determined by grouping those B cells containing identical CDRH3+CDRL3 amino acid sequences. H. Cluster membership separated by clonal expansion for the BM PCs following either LCMV infection or TNFR2 immunization. Color indicates the transcriptional cluster from panel A.

Since we initially observed that TNFR2 immunization and LCMV infection resulted in plasma cells with divergent class-switching profiles (Figures 1D, S1E, S1F), we next questioned whether the sample-specific clustering could be partially explained by isotype-specific gene expression signatures. We therefore overlaid isotype information for each cell based on the antibody repertoire scSeq data, which separated the cells into IgG, IgM, and IgA expressing plasma cells (Figure 3D). Quantifying the cluster membership for all cells of a given isotype demonstrated comparable distributions across the IgG subtypes, which diverged from cells expressing either IgA (cluster 4) or IgM (clusters 4 and 7) (Figure 3E), with exclusive genes defining IgG (*Ly6c2*, *Slpi*), IgA (*Ccl10*, *Glpr1*) and IgM (*Ggh*, *Ptpn6*) (Figure S6). A similar isotype-specific clustering effect was also observed following TNFR2 immunization (Figure 3F), suggesting this effect was not specific to viral infection.

We next asked whether expanded PC clones (clones supported by more than one cell) were transcriptionally distinct compared to unexpanded clones (clones supported by only one single cell). Visualizing and quantifying the cluster membership of the expanded and unexpanded clones following chronic LCMV infection demonstrated similar transcriptional profiles between the two groups (Figures 3G, 3H), implying that either clonal expansion does not result in persisting transcriptional programs in BM PCs or that we have not sampled enough cells to sufficiently label unexpanded cells.

### Clonally expanded plasma cells express antibodies with specificity to viral antigens and have unique gene expression profiles

After observing both high levels of clonal expansion and somatic variants of the IgG isotype in plasma cells of LCMV-infected mice, we next evaluated whether this correlated to antibodies with binding specificity to LCMV antigens. To this end, the scSeq antibody repertoire allowed us to directly synthesize paired V_L_-C_k_-2A-V_H_ cassettes which were cloned into a custom vector for the generation of stable hybridoma cell lines using CRISPR/Cas9 (Parola et al., 2019; Pogson et al., 2016); this way, we recombinantly expressed and validated the specificity for the 30 most expanded IgG clones from the BM PC repertoire (Table S1). We first investigated the specificity of each of the antibodies by ELISA by testing for binding to the following antigens: purified recombinant NP or GPC from LCMV clone 13, lysate from uninfected and LCMV-infected MC57G cells (a C57BL/6-derived fibroblast cell line; Battegay et al., 1991), insulin, dsDNA, and DNP-OVA. Two of the IgG clones displayed clear reactivity against the viral NP protein, and three against GPC, all five of which showed binding to the lysate of infected cells (Figures 4A, 4B, S7A, S7B). Surprisingly, one clone demonstrated reactivity to both the uninfected and infected MC57G lysate (Figures 4A, S7A), potentially indicating the presence of a clonally-expanded BM PC producing autoreactive antibodies. For the three antibody clones that had specificity to GPC, a convergence towards the same variable germline gene combination (IgHV14-4, IgKV10-94) was observed, despite the clones having different CDRH3+CDRL3 amino acid sequences and lengths (Figure 4B). Further investigation into the V gene regions of the three GPC-specific antibodies demonstrated 12 amino acid mutations across either the V_H_ and V_L_ segments despite convergent germline gene usage and CDR3 sequence (Figure S8). Conversely, the NP and uninfected lysate antibodies used distinct germline genes and had more biochemical variation in their CDRH3 regions compared to the GPC binders (Figure 4B). Analyzing the isotype distribution demonstrated IgG subtype heterogeneity across the GPC-specific clones despite the identical germline gene usage and similar CDR3s (Figure 4C). Visualizing these selected clones using UMAP based on their transcriptomes suggested that the NP- and GPC-specific clones occupied transcriptionally distinct states, and quantifying their cluster membership showed they primarily belonged to either cluster 0 or cluster 1, respectively (Figures 4D, E). Differential gene expression analysis revealed that interferon-related genes such as *Ifnar2* and *Ifitm3* were upregulated in cluster 0 whereas genes such as *Tigit*, *CD24a* were upregulated in cluster 1 (Figure S9A). Performing an additional differential expression analysis of all cells binding LCMV revealed that GPC-specific clones significantly upregulated MHC-II genes compared to the NP-specific clones, which in turn demonstrated increased interferon-related genes (Figure S9B). Taken together, these findings support a model in which plasma cells targeting different viral proteins occupy distinct transcriptional states.

**Figure 4.**
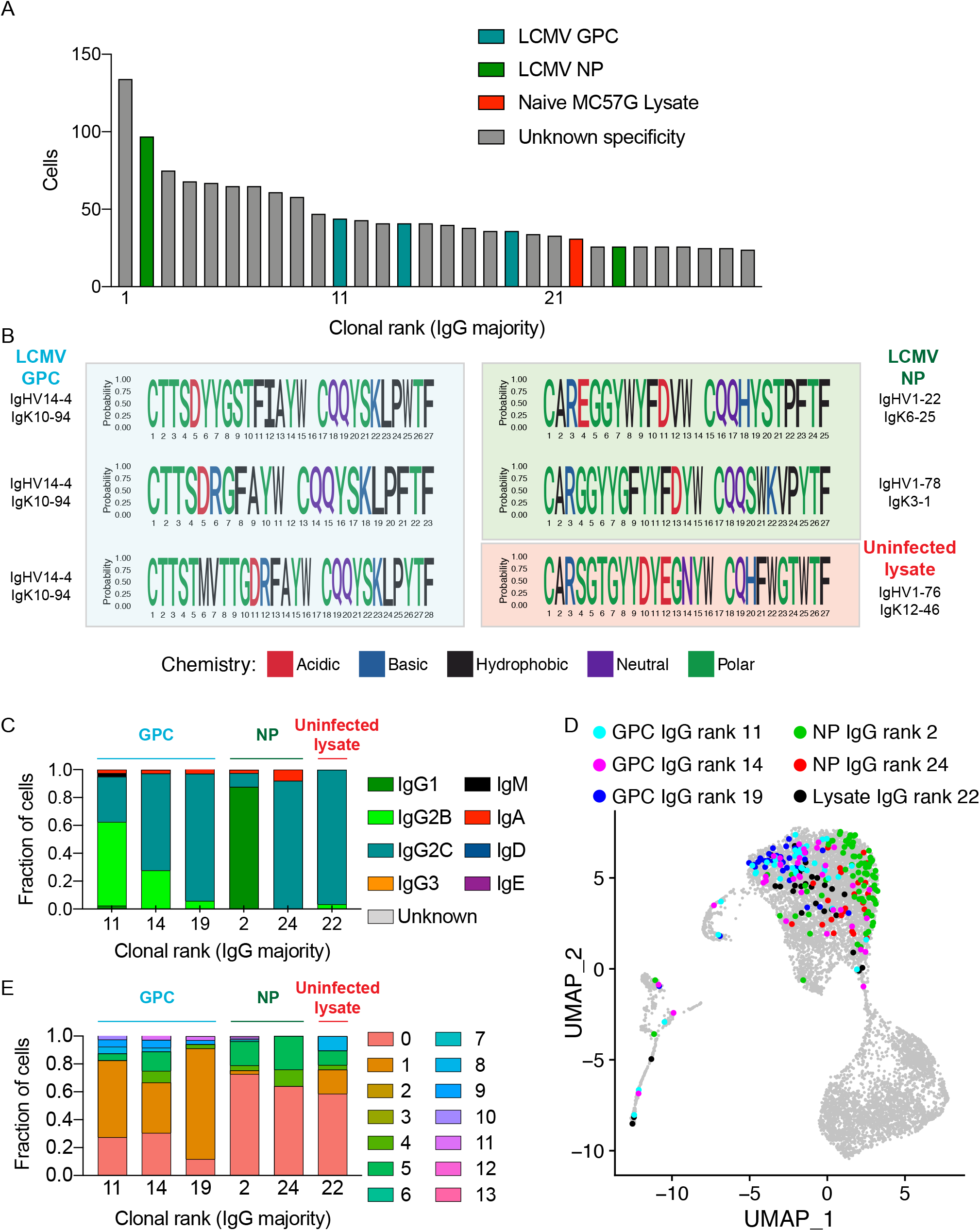
Clonally expanded plasma cells are virus-specific and potentially autoreactive. A. Clonal rank plot of expanded IgG clones indicating viral protein and potentially autoreactive clones. Clone was determined by grouping those B cells containing identical CDRH3+CDRL3 amino acid sequences. Only cells containing exactly one variable heavy (V_H_) and variable light (V_L_) chain were considered. The isotype was determined as the isotype corresponding to the majority of cells within one clone. For each clone, the antibody variant (combined V_H_+V_L_ nucleotide sequence) supported by the most unique cell barcodes was selected to be expressed. B. Sequence logo plots of the confirmed GPC, NP, and MC57G lysate binders. C. Isotype distribution for the confirmed GPC, NP, and MC57G lysate binders. D. Location of the confirmed GPC, NP, and MC57G lysate binders on the UMAP. E. Transcriptome distribution for the confirmed GPC, NP, and MC57G lysate binders. Color indicates the transcriptional cluster from 3A.

### Clonally expanded plasma cells express antibodies with specificity to autoantigens following chronic viral infection

After characterizing the LCMV-specific antibodies, we returned our attention to the clonally expanded IgG variant that demonstrated specificity against lysate of uninfected- and LCMV-infected MC57G cells and determined whether any repertoire and transcriptome signatures of autoreactivity were detectable. We first assigned BM PCs to clonal lineages by grouping cells using identical V and J germline genes for both V_H_ and V_L_ chains, identical CDRH3 and CDRL3 sequence length, and sharing 70% amino acid sequence homology across CDRH3+CDRL3. We then calculated mutational networks by computing distance matrices based on the edit distance and reconstructed mutational histories relative to the unmutated reference germline by successively adding each sequence to the least distant sequence in the network. Investigating the mutational network containing the lysate-binding BM PC demonstrated 17 distinct antibody sequences with varying degrees of clonal expansion, isotype distribution, and transcriptional phenotype (Figures 5A, 5B). Comparing the differentially expressed genes between the potentially autoreactive clone and the virus-specific clones demonstrated 271 and 281 differentially expressed genes (p value < 0.05) compared to BM PCs with GPC- or NP-specific antibodies, although the number of genes was reduced to 3 and 4 following multiple hypothesis correcting, respectively (Figures 5C, 5D). There were no indications of enrichment of genes involved in autoreactivity when performing an unbiased gene ontology analysis (Figures 5E, 5F), which could be expected given the minor transcriptional differences to the virus-specific clones (Figures 5C, 5D). Next we determined whether this clone would demonstrate specificity to murine tissue lysates as it did to the murine MC57G fibroblast cell line lysate. We performed ELISA against lysates from the spleen and bone marrow of uninfected mice. Using both commercial and self-prepared lysate, we could not detect any reactivity relative to the positive control anti-actin antibody (Figure 5G), highlighting the difficulty of discovering the target present in both naive and infected lysate for this clonally expanded IgG antibody.

**Figure 5.**
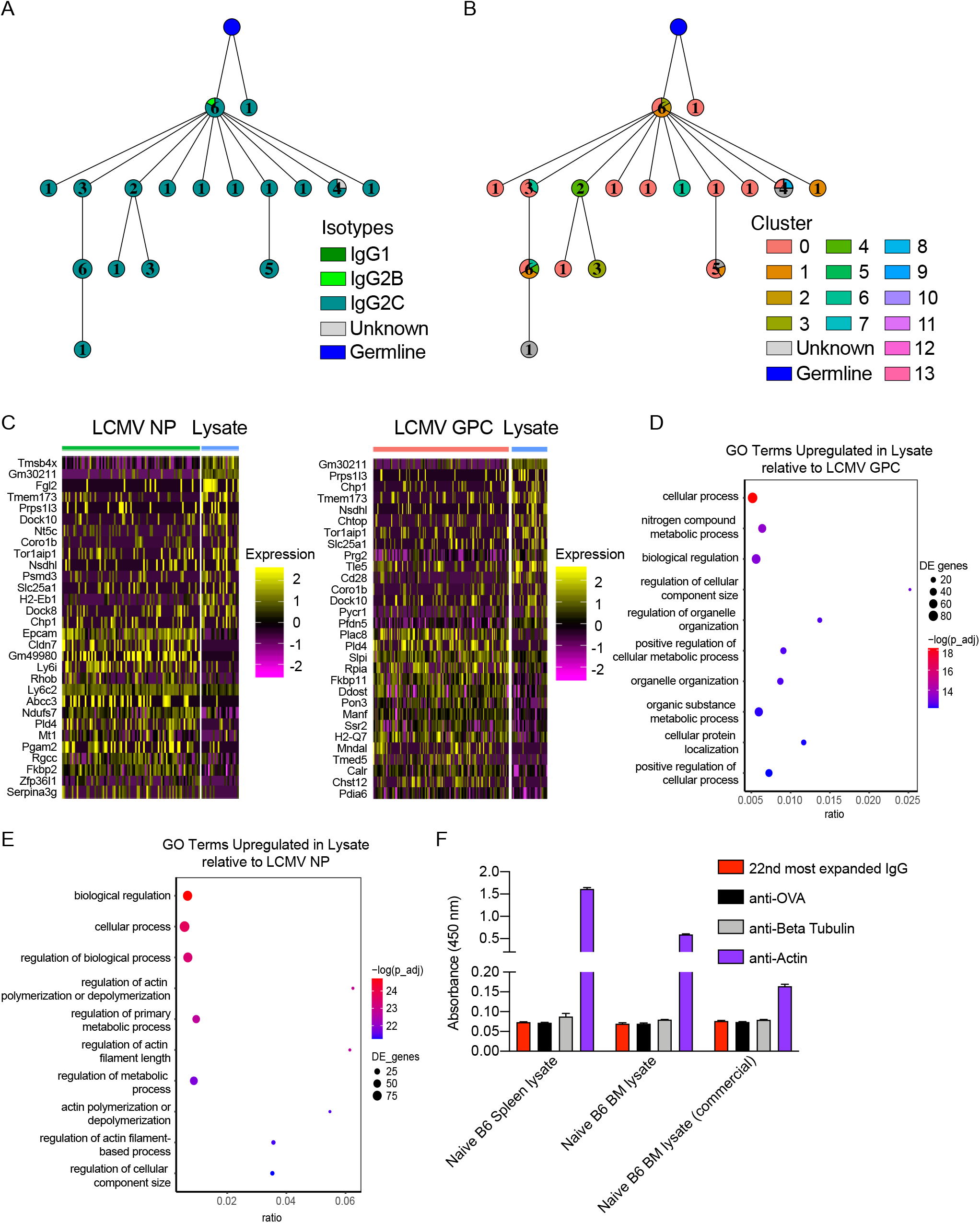
Repertoire and transcriptome profile of MC57G lysate binder. A-B. Mutational network of the IgG clone binding MC57G lysate. Nodes represent unique antibody variants (combined V_H_+V_L_ nucleotide sequence) and edges demonstrate sequences with the smallest separation calculated by edit distance. Node color corresponds to either isotype (A) or transcriptional cluster (B). The size and label of the nodes indicate how many cells express each full-length antibody variant. Clone was determined by grouping those B cells containing identical CDRH3+CDRL3 amino acid sequences. Only cells containing exactly one variable heavy (V_H_) and variable light (V_L_) chain were considered. The isotype was determined as the isotype corresponding to the majority of cells within one clone. The germline node represents the unmutated reference sequence determined by 10X Genomics cellranger. C. Differentially expressed genes between LCMV NP, GPC binders and MC57G lysate binders. The first 15 genes of genes (from top to bottom) corresponds to the highest positive average log fold change (upregulated in MC57G lysate binders). Genes 16-30 represent those genes with the lowest average log fold change (downregulated in MC57G lysate binders). D-E. Gene ontology (GO) term enrichment of upregulated genes in MC57G lysate binder compared to either LCMV GP (D) or LCMV NP (E) binders. The color of each dot corresponds to adjusted p value. The size of the dot corresponds to the number of genes. Ratio corresponds to the number of differentially genes relative to the number of total genes corresponding to each GO term. F. ELISA on lysate from bone marrow (BM) and spleen of naive C57BL/6 mice.

### Virus-specific plasma cells gain cross-reactivity via somatic hypermutation

Since we were unable to determine the specificity of the antibody clone recognizing lysate of uninfected MC57G cells, we shifted our analysis to those clones with confirmed LCMV-specificity. We first analyzed the relationship between somatic hypermutation, clonal expansion, and transcriptional phenotype of specific clonally related antibody variants within virus-specific clonal lineages. Inferring mutational networks for the remaining virus-specific clonal lineages demonstrated heterogeneity across distance of clonally expanded antibody variants from germline, isotype distribution, and transcriptional cluster membership (Figures 6A, 6B, S10, S11). Highlighting the lineage containing the second most expanded clone, which we previously determined as NP-specific (Figure S7), revealed that the most expanded antibody variant (which we had also tested) was closest to the germline, predominantly of the IgG1 isotype, and located primarily in transcriptional cluster 0 (Figures 6A, 6B).

**Figure 6.**
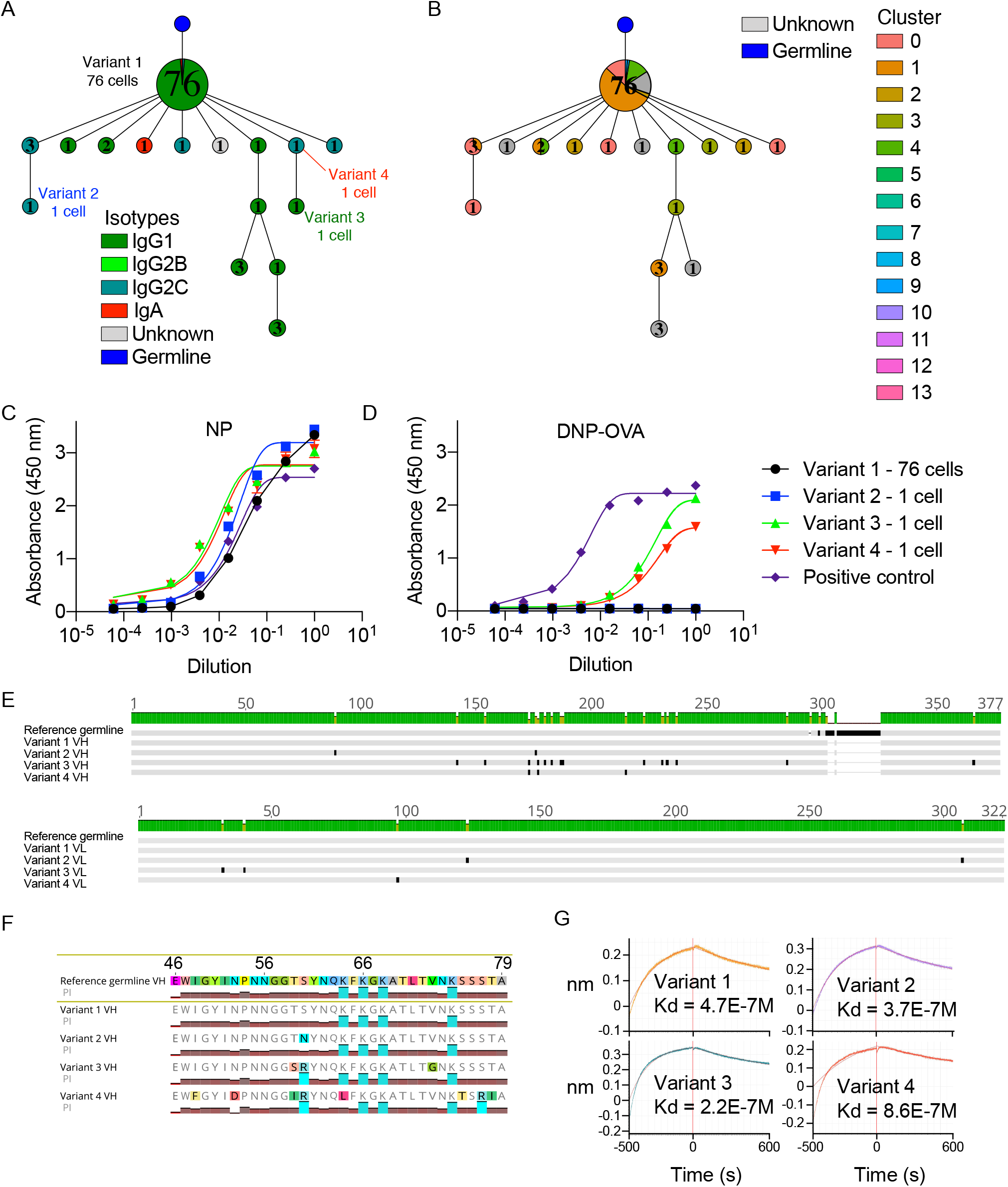
Virus-specific somatic variants present in the BM PC repertoire are cross-reactive. A-B. Mutational network of the NP-specific, second most expanded IgG clone. Nodes represent unique antibody variants (combined V_H_+V_L_ nucleotide sequence) and edges demonstrate sequences with the smallest separation calculated by edit distance. Node color corresponds to either isotype (A) or transcriptional cluster (B). The size and label of the nodes indicate how many cells express each full-length antibody variant. Clone was determined by grouping those B cells containing identical CDRH3+CDRL3 amino acid sequences. Only cells containing exactly one variable heavy (V_H_) and variable light (V_L_) chain were considered. The isotype was determined as the isotype corresponding to the majority of cells within one clone. Variant labels indicate those full-length antibodies which were recombinantly expressed. The germline node represents the unmutated reference sequence determined by 10X Genomics cellranger. C. NP ELISA for the four variants from the second most expanded IgG clone. D. DNP-OVA ELISA for the four variants from the second most expanded IgG clone. E. V_H_ and V_L_ nucleotide alignments of the four variants from the second most expanded IgG clone. F. V_H_ amino acid alignment highlighting mutations in the CDRH2. Colored graphs correspond to isoelectric point (PI). G. Affinity measurements (k_d_) against LCMV NP for the four variants from the second most expanded IgG clone in molar (M) concentrations.

We next addressed whether the mutated variant antibodies produced by unexpanded BM PCs maintained reactivity to NP. We therefore additionally expressed three variants that had incurred somatic hypermutations across both V_H_ and V_L_ chain sequences relative to both the unmutated germline reference sequence and the most expanded BM PC variant in the clonal lineage with confirmed NP specificity (Figures 6C, S7). ELISA confirmed that all three tested variants maintained NP specificity with comparable signals to the previously tested variant and our positive control antibody (Figures 6C, S11A). Surprisingly, when screened for specificity against other antigens (LCMV GPC, insulin, dsDNA, DNP-OVA, lysate of uninfected and infected cells), we discovered that two of the variants specifically bound DNP-OVA but not the other tested LCMV-unrelated antigens (Figures 6D, S12), thereby providing a potential explanation for the LCMV-unspecific IgG antibody response characteristic of chronic viral infection (Greczmiel et al., 2020). Sequence analysis revealed that the three variants had incurred multiple somatic hypermutations, preferentially in their CDRH2 relative to the most expanded variant and reference germline (Figure 6E). Closer inspection revealed that the two cross-reactive variants shared a common S60R mutation in the Vernier region of the CDRH2 relative to the other two single-reactive antibodies (Figure 6F), further supporting the hypothesis that cross-reactivity was acquired by somatic hypermutations and CDRH2 loop flexibility. We lastly determined whether acquiring cross-reactivity reduced the affinity against LCMV NP by measuring the equilibrium dissociation constants (*K*_*d*_) for the four variants. Interestingly, the *K*_*d*_ values were relatively low for all four variants, with no trend that cross-reactivity diminished NP-binding ability (Figure 6G).

## Discussion

Determining the specificity of antibodies produced by plasma cells, residing mainly in the bone marrow, is challenging due to their lack of surface-expression of the BCR (Kräutler et al., 2020; Wolf et al., 2011). Even though there have been significant advances recently to address this problem (Eyer et al., 2017; Gérard et al., 2020; Horns et al., 2020; Setliff et al., 2019), most of previous studies have been reliant upon bulk assays to quantify circulating serum antibodies (ELISA) or antibody secreting cells (ELISpot). These classical approaches fail to inform about the genetics and the clonal diversity of a virus-specific antibody response (i.e., how many unique antibody clones), and about the viral protein specificity of individual BM PCs. Our integrated scSeq approach resolved this short-coming, as we were able to recover thousands of naturally paired V_H_ and V_L_ sequences and their corresponding isotypes, some of which were represented across a multitude of distinct PCs (Figure 1). Overall, we observed differences in isotype distribution, clonal expansion, somatic hypermutation, and gene expression profiles when comparing chronic viral infection to protein immunizations (Figures 1–3, S1). Although these findings need to be interpreted with caution, it nevertheless provided an initial insight into potential differences of identity adopted by BM PCs arising in the context of a chronic viral infection as compared to BM PCs developing after protein immunization.

With the recent advent of scSeq workflows, it is now possible to comprehensively test and reconstruct the specificity of BM PCs based on their clonal expansion profiles. Using high-throughput antibody expression and screening, we were able to demonstrate that a fraction of expanded cells in the bone marrow PC compartment produce antibodies against LCMV-derived NP and GPC antigens following chronic LCMV infection (Figure 4). Given the high abundance of NP protein compared to other viral proteins (King et al., 2018; Pinschewer et al., 2003) in infected cells, it seems intuitive that several of the highest expanded antibody clones were specific against this viral target. We cannot rule out, however, that the viral proteins present at 28 dpi have acquired escape mutations compared to the purified wildtype NP/GPC protein as well as the LCMV clone 13 virus cell lysate that was used for experimental validation, such that antibody binding to these potential variants was not detected in the applied assays. Despite these screening limitations, we were able to demonstrate that during chronic viral infection, somatic hypermutation was implicated in the evolution of viral-specific antibodies into cross-reactive antibodies with specificity to unrelated non-viral antigens (Figure 6). Chronic HIV infection has also been implicated in the development of cross-specificity of HIV-specific antibodies towards other antigens (i.e., phospholipids) (Haynes et al., 2005; Matyas et al., 2009), however the extent of this cross-reactive capacity within a given antigen-specific clonal lineage has not been determined. However, we could further demonstrate in this study that the cross-reactive BM PCs were not clonally expanded within the clonal family (Figure 6), implying that acquiring cross-reactivity does not coincide with a selective advantage, as we would otherwise expect the most expanded antibody variant to demonstrate cross-reactivity. Furthermore, we discovered a clonally expanded IgG class-switched and mutated clone that showed high reactivity to both lysates of uninfected as well as infected MC57G cells (Figures 4, 5) implying autoreactive capacity as has been reported previously for high-dose LCMV infection (Hunziker et al., 2003; Ludewig et al., 2004). Although this cancerous fibroblast cell line originates from a C57BL/6 mouse, we could not demonstrate binding to protein extracts from various tissue lysates from uninfected C57BL/6 mice or other known targets for autoreactive clones such as insulin or dsDNA (Figure S7). In order to determine the binding antigen as well as tissue-specificity, co-immunoprecipitation followed by tandem mass-spectrometry or serological analysis of expression cDNA libraries (SEREX) (Chen et al., 2005; Ludewig et al., 2004) could be used in future studies.

Our data suggests that there may be stereotypical binding preferences for GPC, but not NP reactive clones, as the GPC binding clones utilized identical germline gene combinations and similar CDR3 sequences (Figure 4) (Hsiao et al., 2020). In contrast, the NP specific BM PCs utilized different germline genes (Figure 4), consistent with previous V_H_ repertoire sequencing results, which demonstrated a continuous recruitment of unique clones into the BM PC repertoire over time during chronic but not acute LCMV infection (Kräutler et al., 2020). Although our study relies on the bone marrow PC compartment of a single infected mouse, we have nevertheless demonstrated that our scSeq pipeline can be useful in profiling a personalized, virus-specific plasma cell response.

Finally, our study linked antibody repertoire information with total transcriptome profiles, providing a detailed transcriptional description of distinct plasma cell clonotypes in the bone marrow. Our results suggest that IgM- and IgA-secreting BM PCs have distinct transcriptional profiles compared to the IgG clones, and that within virus-specific IgG clones there was transcriptional diversity (Figure 4). Some antibody discovery pipelines rely heavily on class-switched IgG memory B cells following immunization schemes (Parola et al., 2018), leveraging the ability to directly stain surface BCRs and identify antigen-specific B cells via flow cytometry. The lack of surface BCR/antibody on BM PCs means that a scSeq approach was required to interrogate binding specificity. This also allowed us to pair transcriptome information with antibody sequence which further revealed that BM PCs producing either LCMV NP- or GPC-specific antibodies had distinct transcriptional profiles (Figure 4). While this analysis was restricted to approximately 100 cells for both antigens, it nevertheless demonstrated an increase of MHC-II gene expression in the GPC-specific BM PCs, in contrast to an increase of inflammatory interferon-induced genes in the NP-specific BM PCs (Figure 4). This transcriptional program may reflect distinct selection pressures for GPC- in comparison to NP-specific BM PCs at this time point, thereby promoting the onset of neutralizing antibodies. Future studies analyzing GPC and NP-specific BM PCs at early and later time points could better delineate whether such a transcriptional phenotype is exclusively reserved for GPC-specific BM PCs, or reflects longitudinal GC selection based on which viral protein complexes are displayed and recognized on follicular dendritic cells.

## Methods

### Mouse experiments

All animal experiments were performed in accordance with institutional guidelines and Swiss federal regulations. Experiments were approved by the veterinary office of the canton of Zurich (animal experimentation permissions 115/2017) and Basel-Stadt (animal experimentation permission 2582). A 6 week old C57BL/6 mouse was infected with 2×10^6^ LCMV clone 13 i.v. as previously described (Kräutler et al., 2020) and sacrificed 28 dpi. LCMV clone 13 was propagated by infection of BHK-21 cells for 24-48 hours and then resuspended in sterile PBS (GIBCO).

For protein immunizations, a 6-week-old C57BL/6 mouse was repeatedly immunized 5 times every other week subcutaneously (s.c.) into the flank with 10ug of human TNFR2 protein (Peprotech, 310-12)/20ug MPLA (Sigma, L6895) adjuvant and sacrificed one week afterwards. Likewise, a 6-week old BALB/c mouse was injected three times with four and three week intervals s.c. into the flank with 100ug of OVA (Sigma, A5503)/20ug MPLA (Sigma, L6895) adjuvant and sacrificed two weeks after the final boost.

### Isolation of bone marrow plasma cells

A single cell suspension was prepared by flushing the bone marrow from the tibia, femur, and hip bones in RPMI containing 10% FCS buffer with 10 ng/mL IL6 (Peprotech, 216-16). A red blood cell lysis step was performed in 4 mL ACK lysis buffer for 1 minute at room temperature and subsequently inactivated with 26 mL RPMI containing 10% FCS. For LCMV infection and TNFR2 immunizations, single-cell suspensions were stained with the following antibodies (1:200 dilution) CD138-APC, CD4-APC-Cy7, CD8a-APC-Cy7, NK1.1-APCCy7, Ter119-APC-Cy7, TACI-PE, B220-APC, CD19-PE-Cy7 for 30 minutes at 4°C. Cell sorting was performed using a FACSAria with FACSDiva software into RPMI. For the OVA immunized mouse, BM PCs were isolated using a CD138+ plasma cell isolation kit (Miltenyi, 130-092-530) following the instructions of the manufacturer.

### Library construction and sequencing

Single cell 10X VDJ libraries were constructed from the isolated bone marrow plasma cells following the demonstrated protocol ‘Direct target enrichment - Chromium Single Cell V(D)J Reagent Kits’ (CG000166 REV A). Briefly, single cells were co-encapsulated with gel beads (10X Genomics, 1000006) in droplets using 3 lanes of one Chromium Single Cell A Chip (10X Genomics, 1000009) with a target loading of 13,000 cells per reaction. V(D)J library construction was carried out using the Chromium Single Cell 5’ Library Kit (10X Genomics, 1000006) and the Chromium Single Cell V(D)J Enrichment Kit, Mouse B Cell (10X Genomics, 1000072). Final libraries were pooled and sequenced on the Illumina NextSeq 500 platform (mid output, 300 cycles, paired-end reads) using an input concentration of 1.8 pM with 5% PhiX.

### Repertoire and transcriptome analysis

Paired-end raw sequencing files from Illumina NextSeq500 run were aligned to murine reference genome and germlines (GRCm38) using 10X Genomics cellranger with clonotyping based on those B cells containing identical CDRH3+CDRL3 nucleotide sequences. Gene expression analysis was performed using the R package Seurat (Satija et al., 2015) using the default parameters from the R package Platypus (Yermanos et al., 2020). Those cells lacking full-length, paired (V_H_+V_L_) antibody sequences in the VDJ library were removed from the transcriptome analysis. Furthermore, genes dictating clonal diversity (e.g. IgH V genes, constant genes, IgH J genes) were removed from the gene expression matrix where indicated. Cells containing more than 5% mitochondrial genes were filtered out before log-normalization using a scaling factor of 10000. Mean expression and variance were further scaled to 0 and 1, respectively. Graph-based clustering incorporating Louvain modularity optimization and hierarchical clustering was performed by the Seurat functions FindNeighbors and FindClusters (Satija et al., 2015). When calculating clonal frequencies, those clones sharing identical CDRH3+CDRL3 amino acid sequences were clustered into a single clonal family. Full-length sequences ranging from framework region 1 to framework region 4 were annotated using the built-in murine reference alleles in MiXCR and exported by the VDJRegion gene feature (v3.0.1). The full-length, heavy and light VDJRegion sequences corresponding to identical cellular barcodes were appended and then the number of clonal variants was determined by calculating the number of unique amino acid sequences. These full-length, appended V_H_+V_L_ sequences were subsequently used to create mutational networks, which were constructed by first clustering all B cells containing identical V and J segments for heavy and light chain, identical CDRH3 and CDRL3 sequence lengths, and sharing 70% sequence homology. The pairwise edit distance was subsequently calculated for each full-length VDJ sequence (including the reference germline), generating a distance matrix to determine the order in which sequences were added to the network. The unmutated germline reference gene (determined by cellranger alignments) initialized each network, and then the sequence with the smallest distance was added to the most similar sequence in the network in an iterative manner. Edit distance ties were resolved by randomly sampling all possible nodes. The final adjacency matrix was visualized as a graph by using the graph_from_adjacency_matrix function in the R package igraph (Csardi et al., 2006). Cluster-specific genes were determined by Seurat’s FindMarkers function. Differential gene analysis for all comparisons was performed using the FindMarkers function in Seurat under default parameters. Cell isotype for gene expression data was extracted by matching barcodes from the VDJ sequencing data. Multiple string alignments and visualizations were performed using Geneious. Gene ontology analysis was performing using the function goanna in the R package edgeR under default parameters (Robinson et al., 2010).

### Antibody expression and validation

Mouse antibodies were produced as previously described (Parola et al., 2019) and validated using normalized supernatant ELISAs against purified NP, GPC and DNP-OVA, insulin (Sigma, I5500), mouse genomic DNA (Sigma, 692339) as well as in house-produced lysate of (LCMV-infected) MC57G cells as previously described (Greczmiel et al., 2017). Recombinant LCMV clone 13 NP and GPC protein were expressed as previously described (Greczmiel et al., 2017; Hastie et al., 2017). An anti-mouse IgG-HRP (Sigma, A2554) was employed at 1:1500 and used for detection. In order to test for tissue specificity of the autoreactive clone, protein extracts from commercial bone marrow tissue (Zyagen, MT-704-C57) as well as self-made extracts from spleen and bone marrow were tested at a 100ug/ml coating concentration. Self-made extracts were prepared from spleen and bone marrow single-cell suspensions. Cells were centrifuged 5 min at 1600 rpm at 4°C and the pellet was resuspended in 5 ml of PBS. Cells were lysed using a syringe (28G needle), homogenizing several times and additionally sonicated (3 times 20 s at 40 MHz) on ice. Anti-HEL antibody (in house) as well as anti-mouse beta tubulin IgG (Sigma, T5201), anti-mouse beta actin IgG (Sigma, A2228), anti-OVA IgG (in house), anti-NP-IgG Pank1, anti-GPC-IgG Wen1.3, anti-insulin IgG E11D7 (Sigma, 05-1066) and anti-dsDNA IgG AE-2 (Sigma, MAB1293) were used as negative and positive controls for respective experiments and employed at 4ug/ml.

### Antibody affinity measurements

Antibody affinity measurements were carried out as reported previously (Mason et al., 2019; Parola et al., 2019). In brief, biolayer interferometry assays were performed on the Octet Red96 (ForteBio) at 25°C, shaking at 1000 rpm. Kinetics assays were performed with Anti-Mouse IgG Fc Capture (AMC) Biosensors (ForteBio, Cat-No. 18-5090) with the following steps: (0) Hydration of AMC Biosensors in 1X kinetics buffer for 30 mins (ForteBio, Cat-No. 18-1105). (1) Baseline equilibration in conditioned medium diluted 1:1 with 1X kinetics buffer for 60s. (2) Regeneration of sensors (3x) in 10 mM Glycine. (3) Baseline for 300s. (4) Loading of antibodies contained in supernatant for 500s. (5) Quenching/Blocking of sensors in 50 μg/mL hIgG in 1X KB for 30s (6) Antigen association: sensors immersed with NP at 3.2 – 2000 nM for 500s. (7) Dissociation in 1X KB for 600s. (8) Regeneration of sensors. Curve fitting was performed using the ForteBio Octet HTX data analysis software using a 1:1 model, and a baseline correction using a reference sensor.

## Data visualization

Heatmaps displaying differential gene expression were produced using the DoHeatmap function in the R package Seurat (Butler et al., 2018). Gene enrichment plots were produced using the R package ggplot (Wickham and Wickham, 2007). Mutational networks were produced using the R package igraph (Csardi et al., 2006). Sequence logo plots were generated using the R package ggseqlogo (Wagih, 2017). Sequence alignment plots were exported from Geneious Prime. All other figures were produced using Prism v9 (Graphpad), excluding the graphical abstract, which was created with BioRender.com.

## Acknowledgements

We acknowledge and thank Dr. Christian Beisel, Elodie Burcklen, Ina Nissan, and Tobias Schär at the ETH Zurich D-BSSE Genomics Facility Basel for excellent support and assistance. We also thank Mariangela Di Tacchio and Marie-Didiée Hussherr for excellent experimental support.

## Supporting information

**Figure S1.**
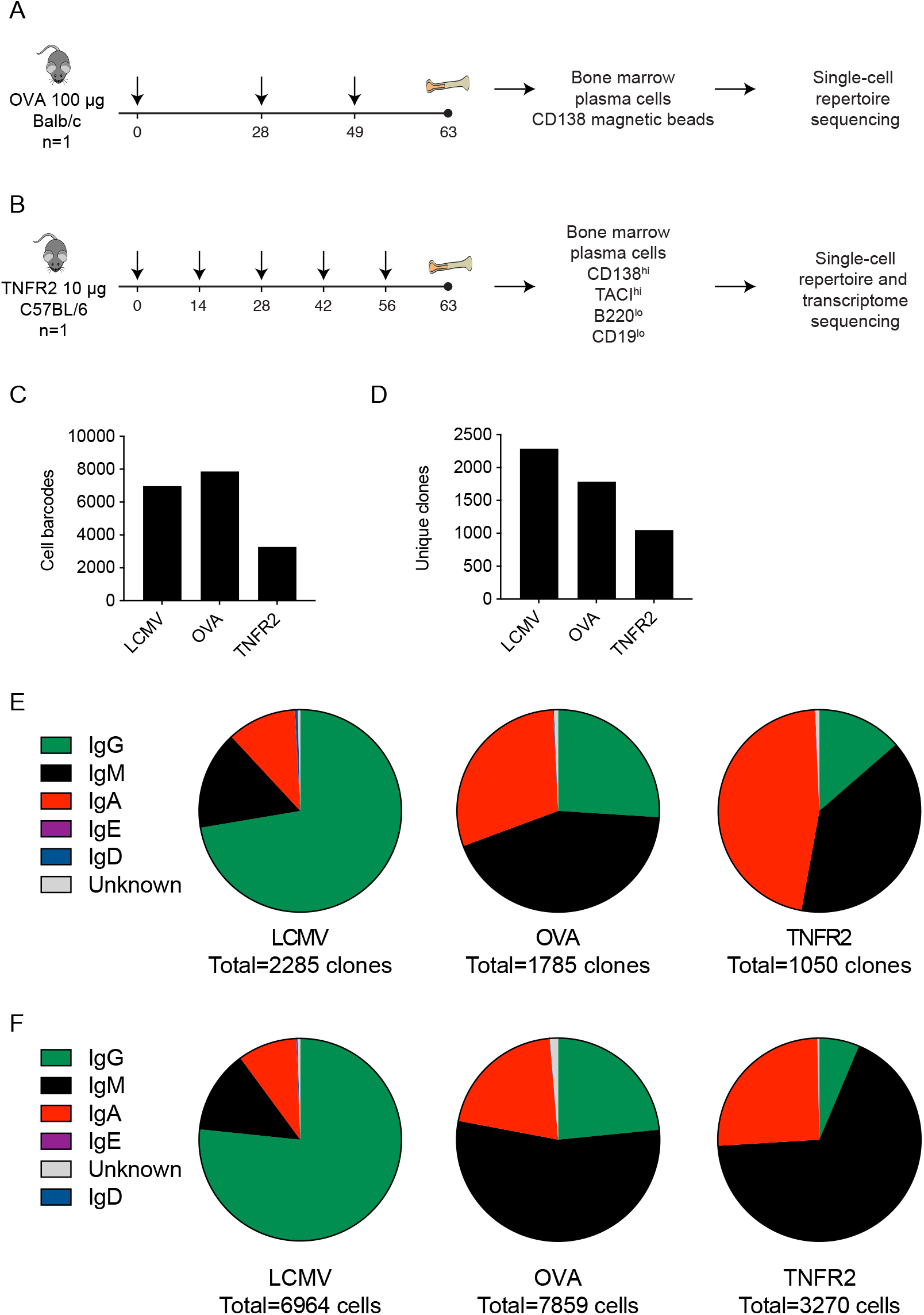
Experimental overview of immunization schemes and antibody repertoire statistics. A. Serial immunization scheme with ovalbumin (OVA) and subsequent bone marrow plasma cell (BM PC) isolation using CD138+ magnetic beads. Subsequent antibody repertoire libraries were prepared using the 5’ single-cell immune profiling kit from 10X Genomics. B. Serial immunization scheme with the extracellular region of human tumor necrosis factor receptor 2 (TNFR2) and subsequent bone marrow plasma cell (BM PC) isolation using flow cytometry. BM PCs were sorted based on CD138^hi^, TACI^hi^, B220^lo^, CD19^lo^. Subsequent antibody repertoire and transcriptome libraries were prepared using the 5’ single-cell immune profiling kit from 10X Genomics. C. The total number of cells recovered in each infected or immunized mouse. Only cells containing exactly one variable heavy (V_H_) and variable light (V_L_) chain were considered. D. The total number of clones recovered in each infected or immunized mouse. Clone was determined by grouping those B cells containing identical CDRH3+CDRL3 amino acid sequences. Only cells containing exactly one variable heavy (V_H_) and variable light (V_L_) chain were considered. E-F. Fraction of clones (E) and cells (F) corresponding to a given isotype in each infected or immunized mouse. Clone was determined by grouping those B cells containing identical CDRH3+CDRL3 amino acid sequences. Only cells containing exactly one variable heavy (V_H_) and variable light (V_L_) chain were considered.

**Figure S2.**
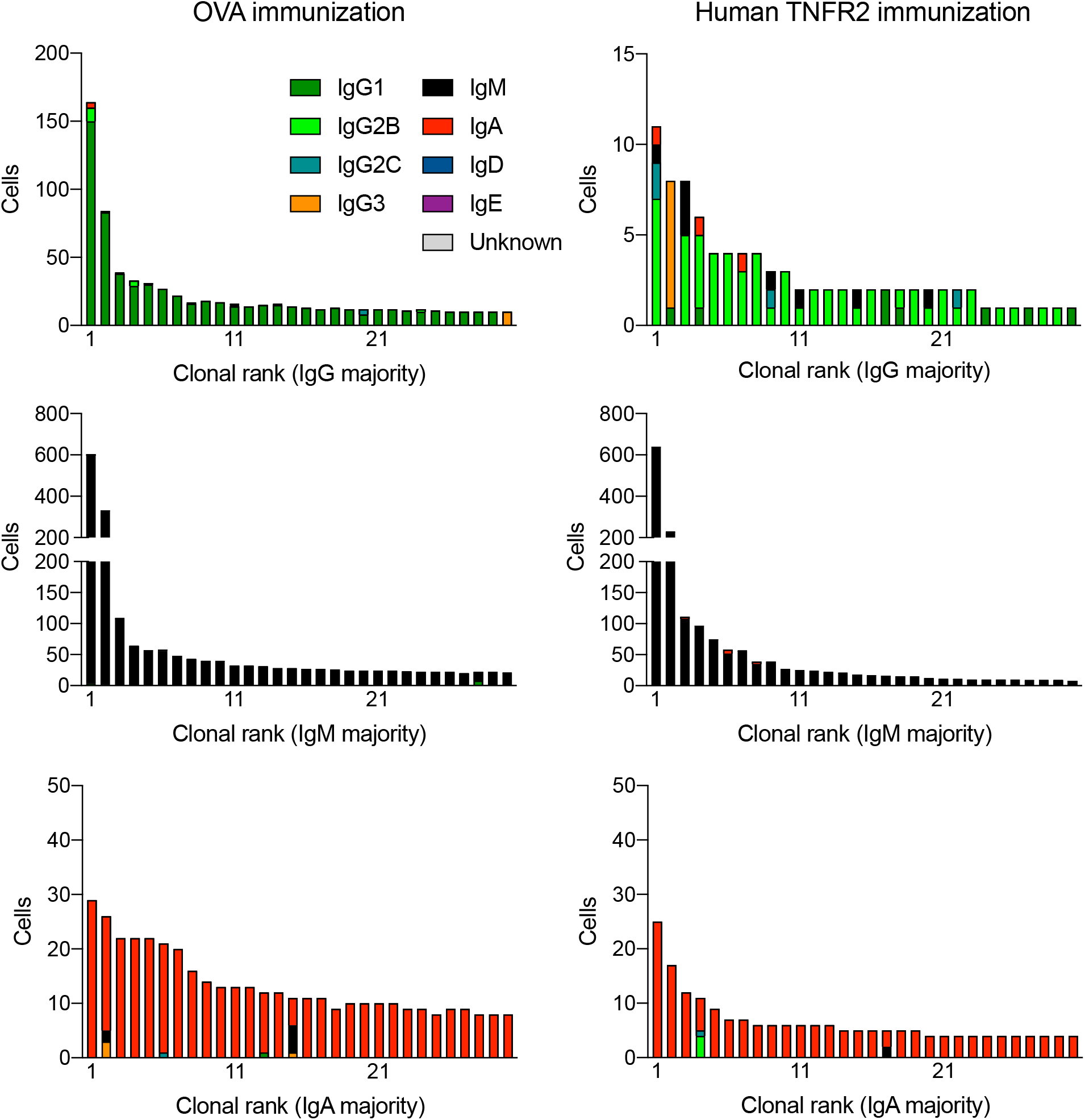
Clonal expansion of bone marrow plasma cells (BM PC) separated by isotype for OVA and TNFR2 immunized mice. The number of distinct cell barcodes belonging to the top 30 clones is visualized. Clone was determined by grouping those B cells containing identical CDRH3+CDRL3 amino acid sequences. Only cells containing exactly one variable heavy (V_H_) and variable light (V_L_) chain were considered. Color corresponds to isotype per each cell.

**Figure S3.**
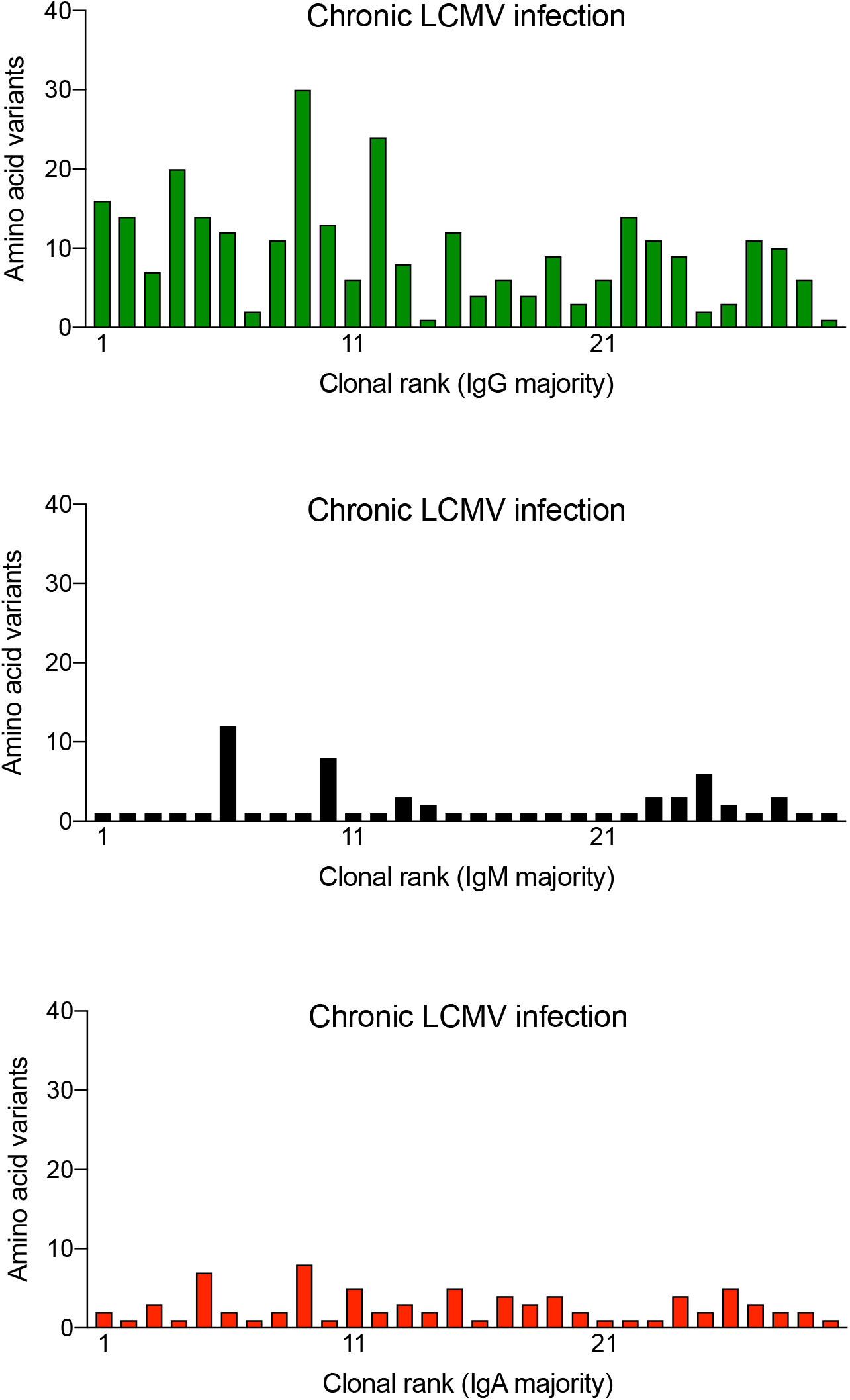
Number of full-length (paired V_H_+V_L_) amino acid variants for the 30 most expanded clones separated by isotype in the bone marrow plasma cell repertoire following chronic LCMV infection. Clone was determined by grouping those B cells containing identical CDRH3+CDRL3 amino acid sequences.

**Figure S4.**
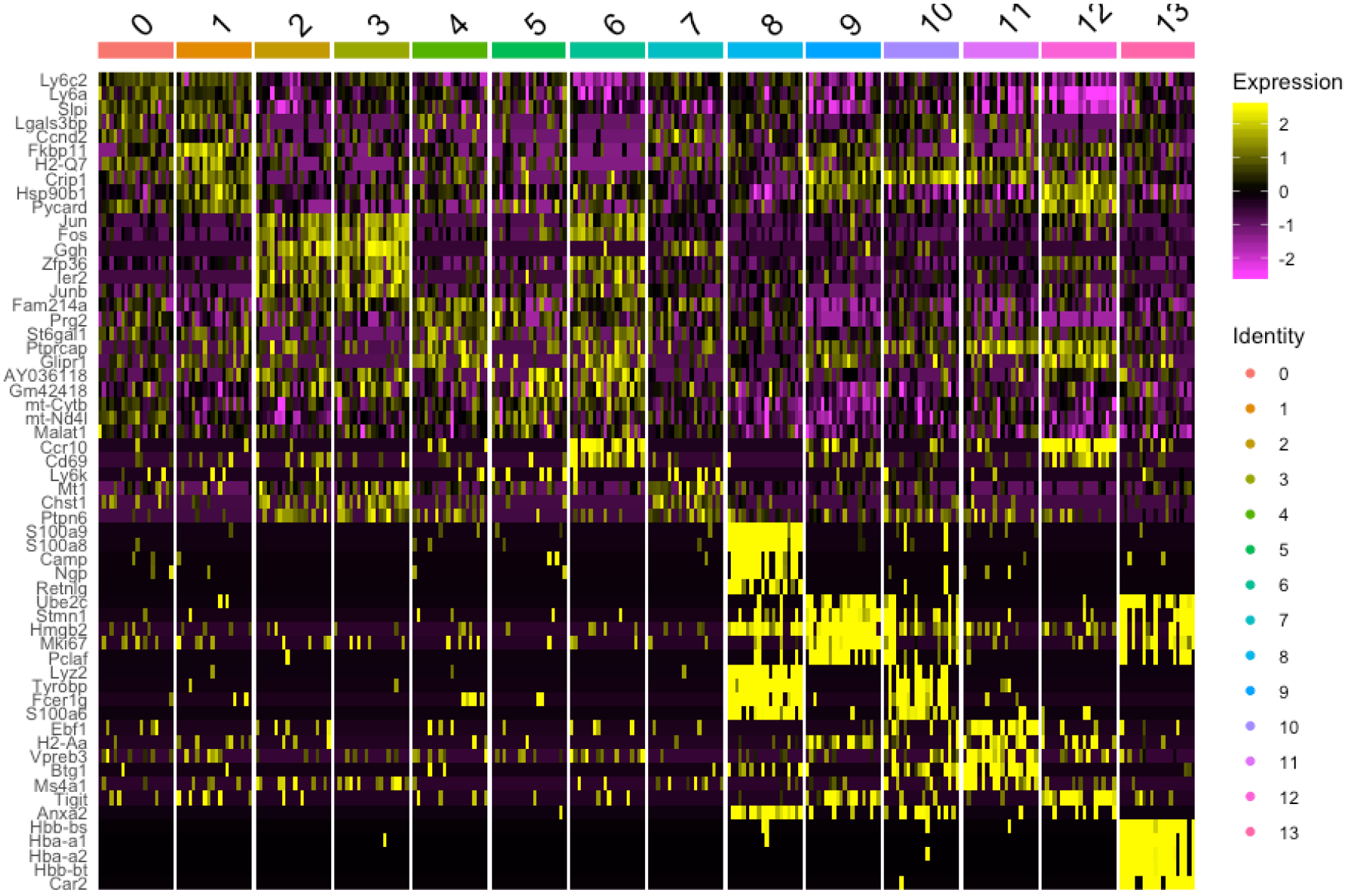
Top differentially expressed genes for each transcriptional cluster of the bone marrow plasma cells from mice either chronically infected with LCMV or immunized with TNFR2. Heatmap intensity corresponds to normalized expression. Each column represents a single cell and each row corresponds to a single gene. The top five genes based on average log fold change (logFC) have been selected for each cluster after removing those ribosomal protein L (RPL), ribosomal protein S (RPS), and mitochondrial genes. All displayed genes had an adjusted p value less than or equal to 0.01.

**Figure S5.**
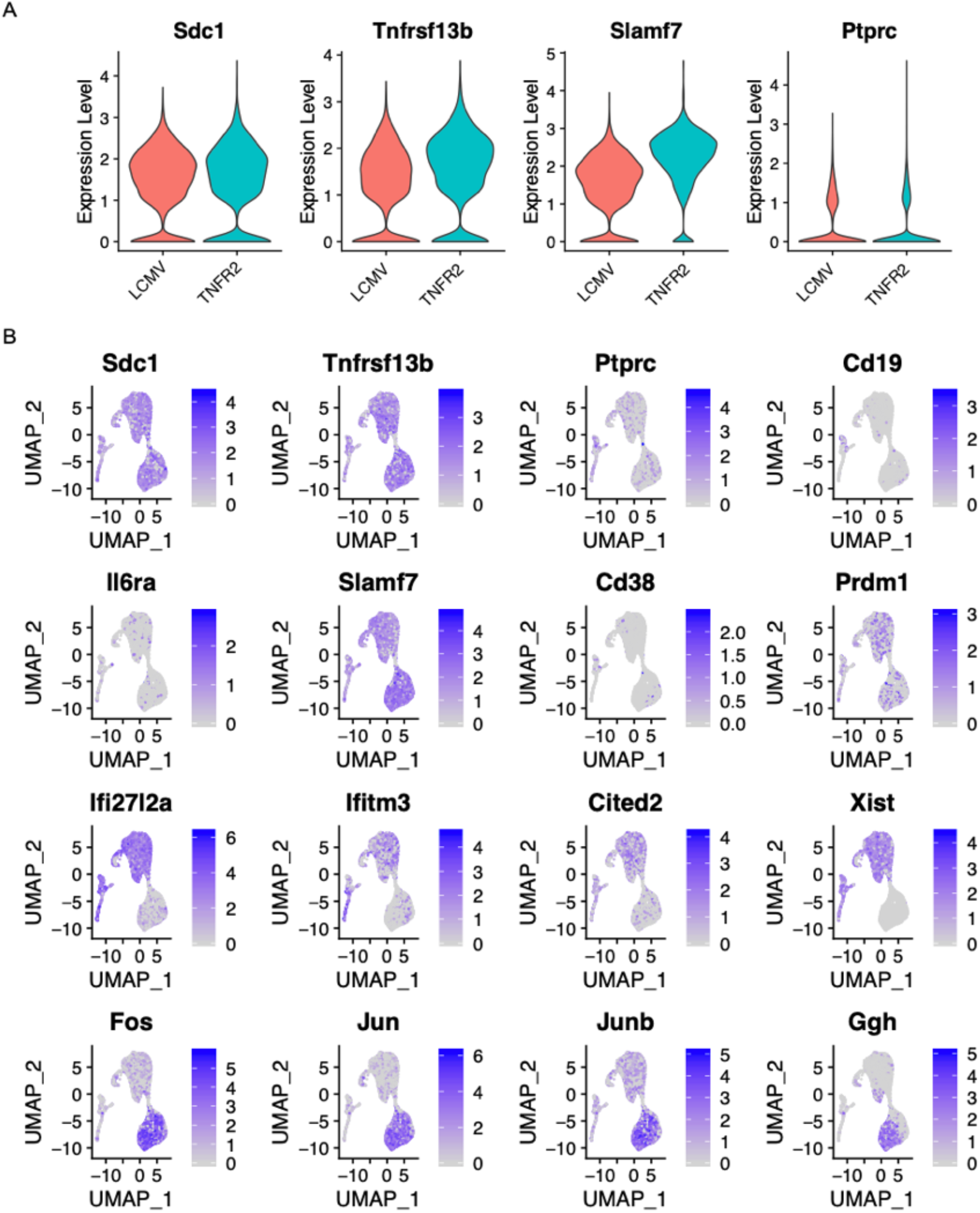
Differentially expressed genes between the bone marrow plasma cells from mice either chronically infected with LCMV or immunized with human TNFR2. (A) Normalized expression of plasma cell genes *Sdc1* (*CD138*), *Tnfrsf13b* (*TACI*), *Slamf7*, and *Ptprc* (*B220*) for either mice infected with LCMV or immunized with human TNFR2. (B) Uniform manifold approximation project (UMAP) plots showing normalized gene expression for selected genes.

**Figure S6.**
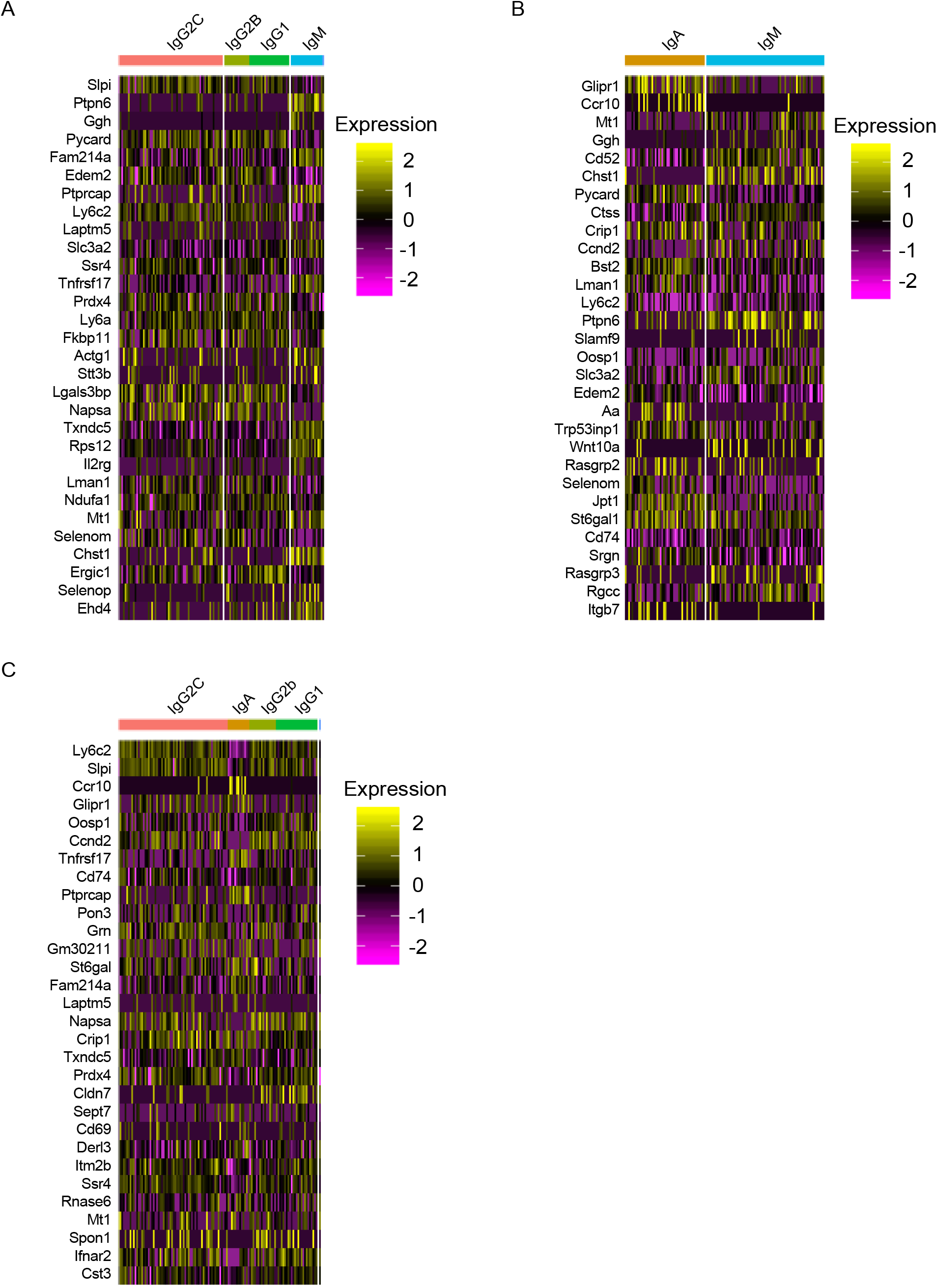
Differentially expressed genes between cells expressing different isotypes. (A) Differentially expressed genes between IgG vs IgM of plasma cells coming from an LCMV infected mouse. (B) Differentially expressed genes between IgM vs IgA of plasma cells coming from an LCMV infected mouse. (C) Differentially expressed genes between IgG vs IgA of plasma cells coming from an LCMV infected mouse. Heatmap intensity corresponds to normalized expression. Each column represents a single cell and each row corresponds to a single gene. The top 30 genes based on average log fold change (logFC) for each isotype comparison were selected. All displayed genes had an adjusted p value less than or equal to 0.01.

**Figure S7.**
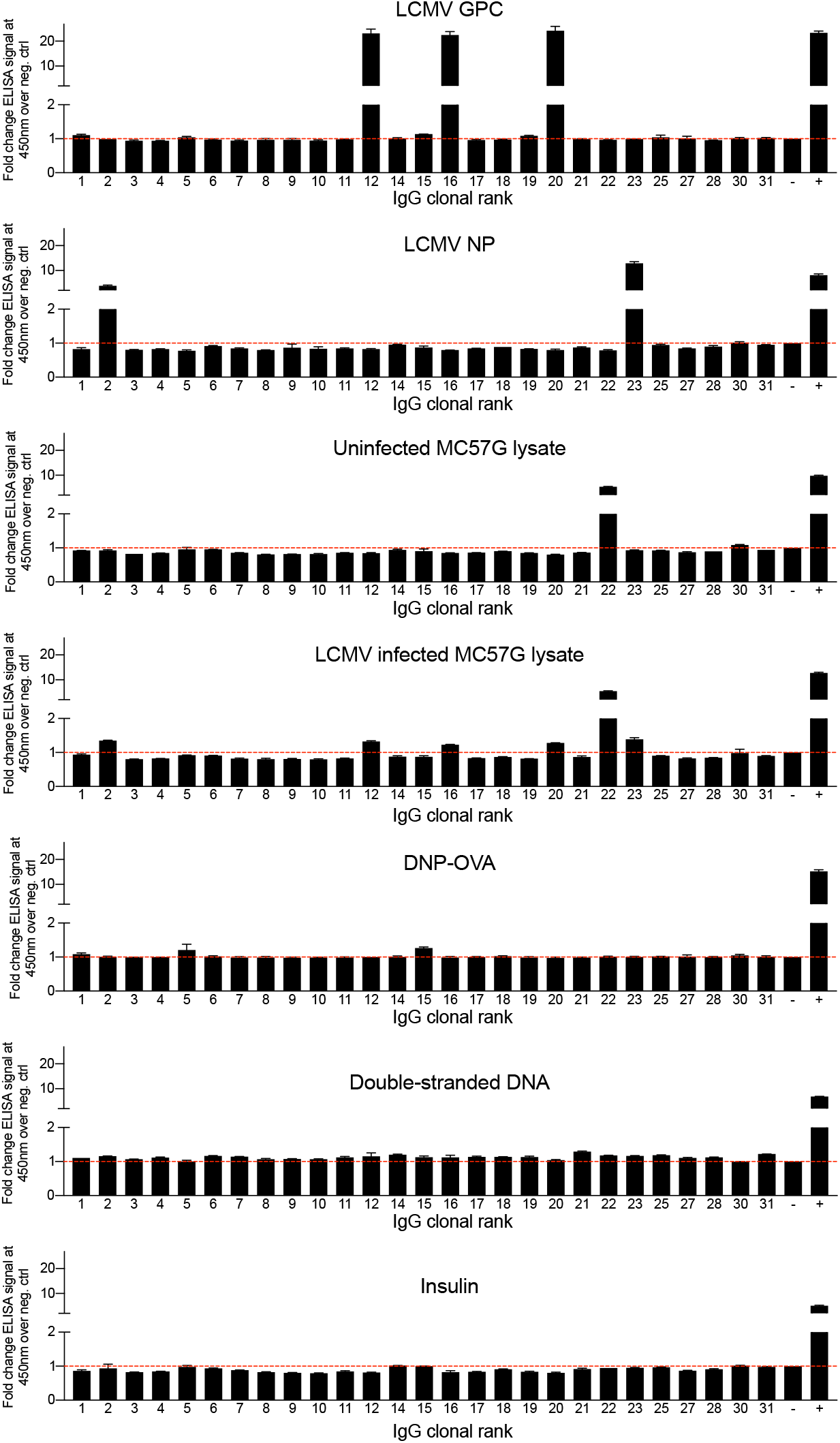
Clonally expanded plasma cells produce virus-specific and potentially autoreactive antibodies following chronic viral infection. The ELISA signal of duplicate ELISA measurements at 450 nm is shown relative to a negative background control (red dotted line indicates background level). Control antibodies used are listed in the methods section.

**Figure S8.**
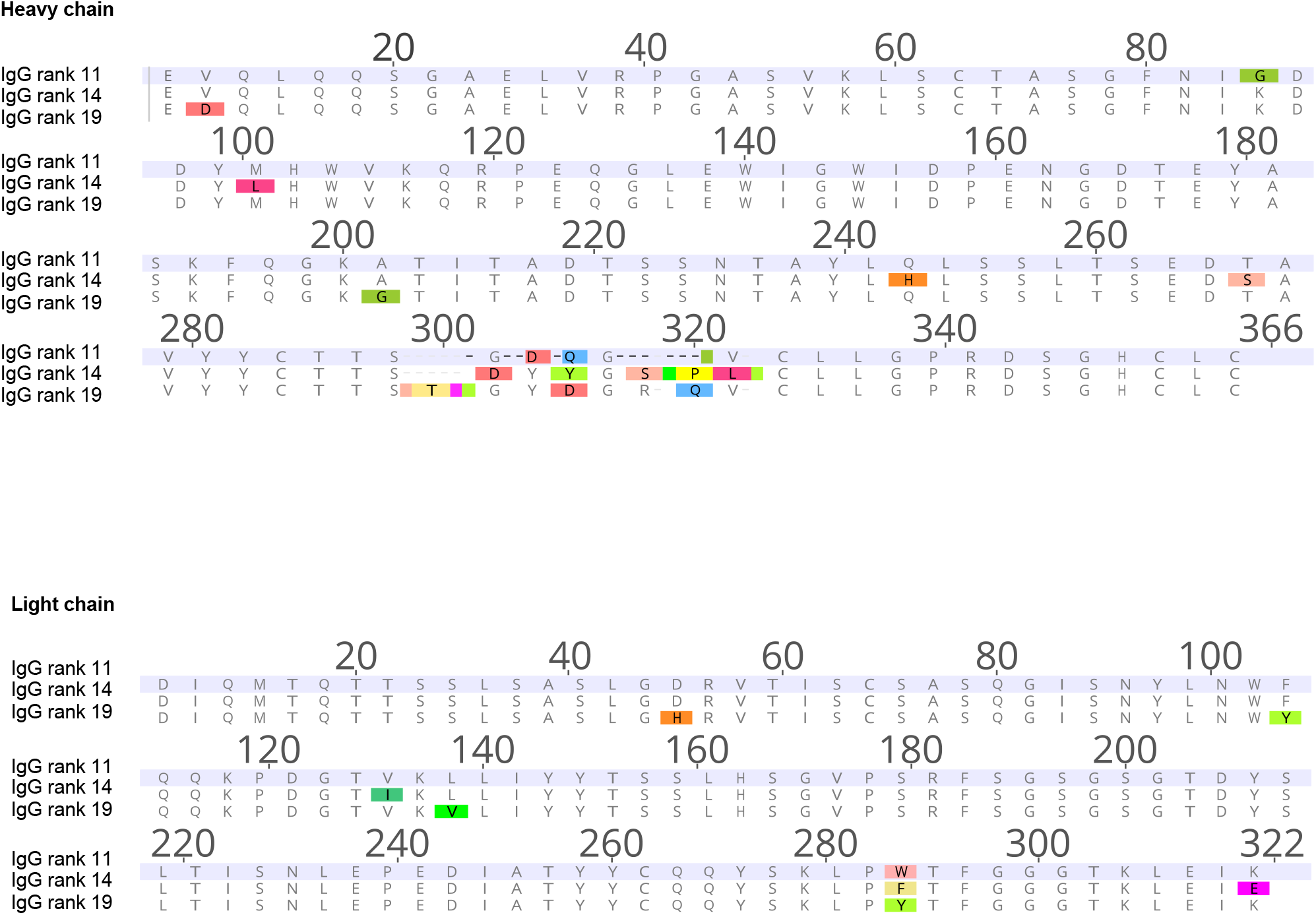
V_H_ and V_L_ amino acid alignment of the three LCMV GPC specific clones.

**Figure S9.**
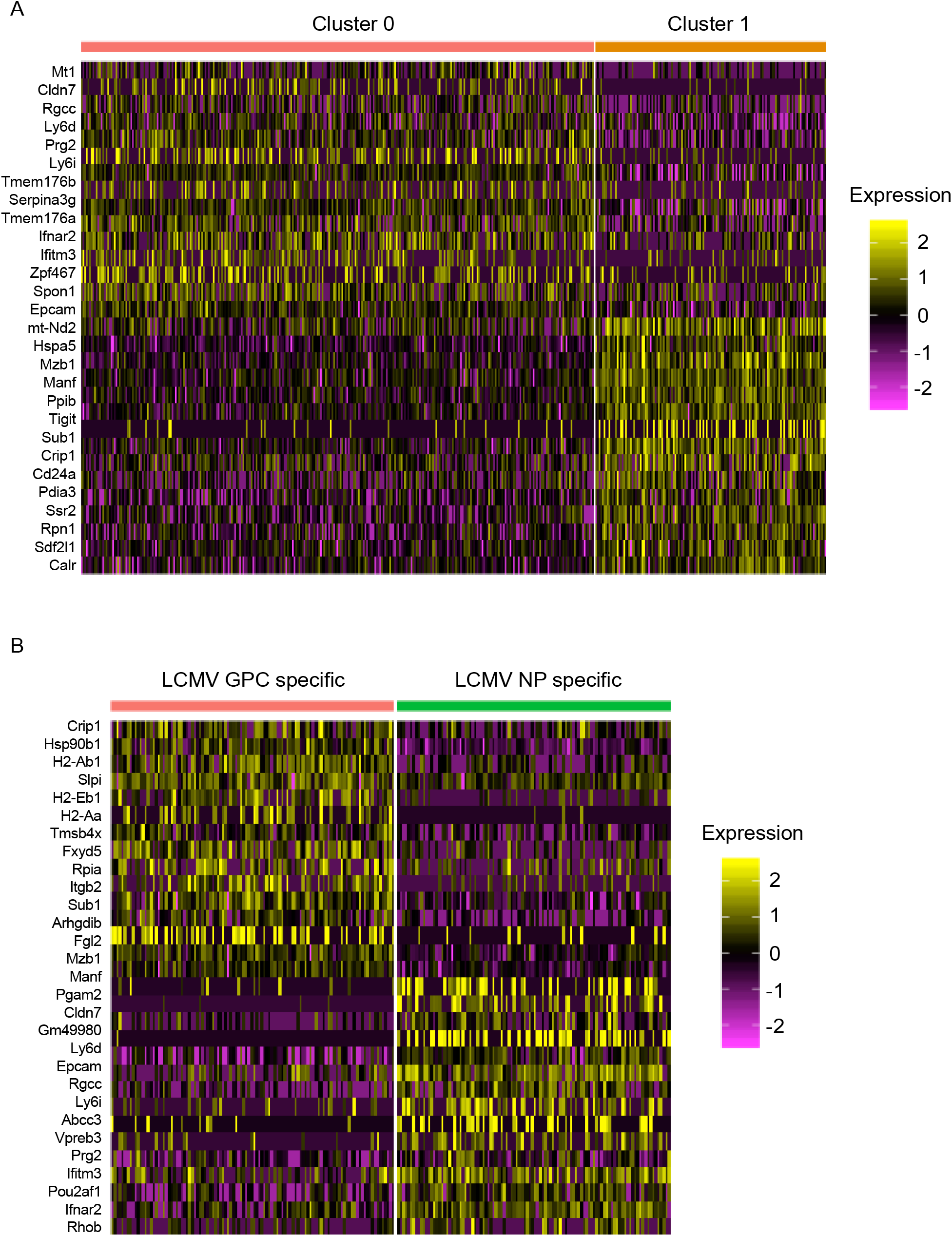
Signature genes expressed by plasma cells producing either NP or GPC specific antibodies. A. Differentially expressed genes between cells located in either cluster 0 or cluster 1. Heatmap intensity corresponds to normalized expression. Each column represents a single cell and each row corresponds to a single gene. The top 40 differentially expressed genes based on average log fold change (logFC) were selected. All displayed genes had an adjusted p value less than or equal to 0.01. B. Differentially expressed genes between the plasma cells producing either NP or GPC specific antibodies. Each column represents a single cell and each row corresponds to a single gene. The top 40 differentially expressed genes based on average logFC were selected. All displayed genes had an adjusted p value less than or equal to 0.01.

**Figure S10.**
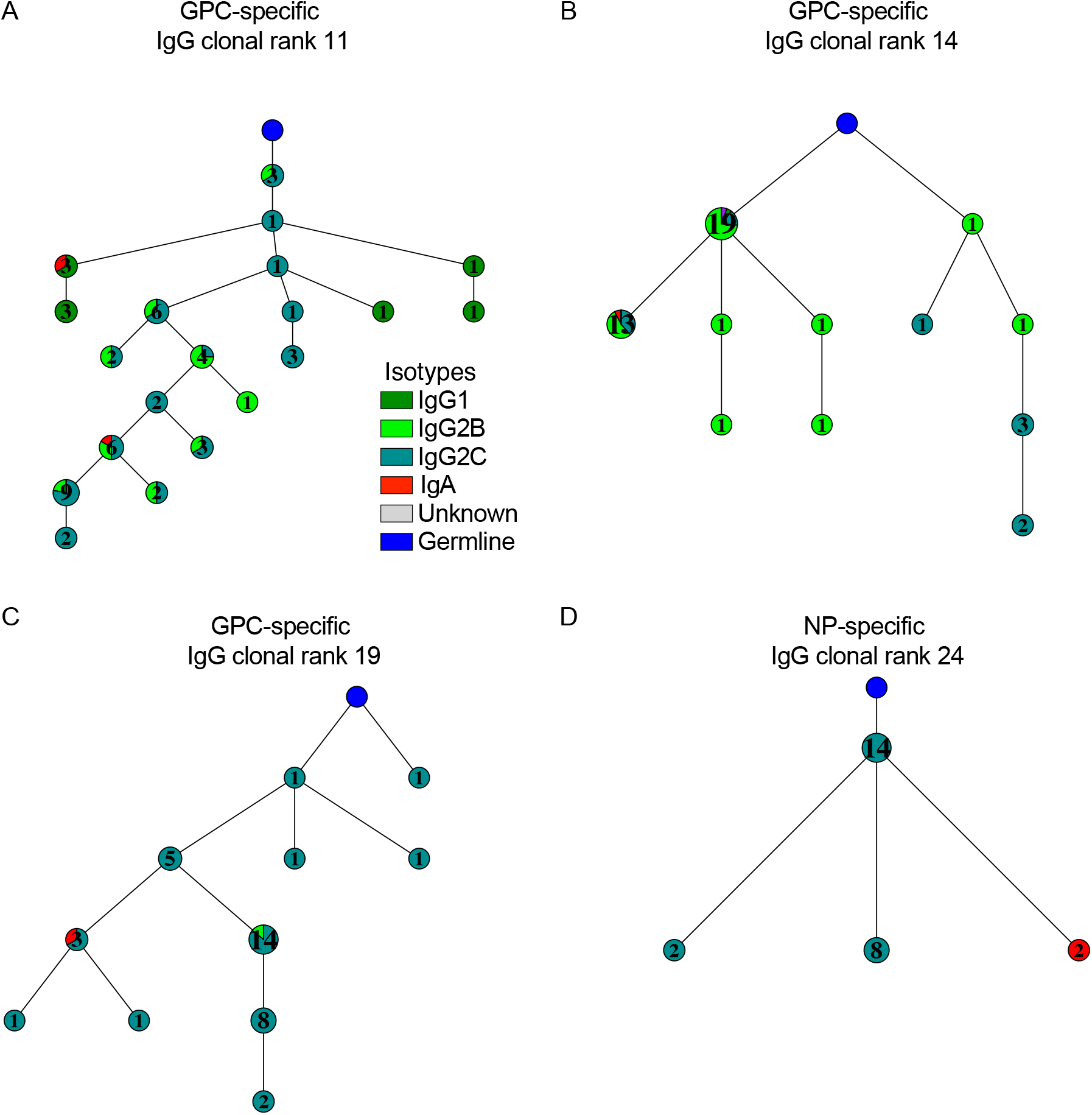
Virus-specific mutational networks displaying isotype distribution. Nodes represent unique antibody variants (combined V_H_+V_L_ nucleotide sequence) and edges demonstrate sequences with the smallest separation calculated by edit distance. Node color corresponds to isotype distribution for each cell. The size and label of the nodes indicate how many cells express each full-length antibody variant. Clone was determined by grouping those B cells containing identical CDRH3+CDRL3 amino acid sequences. Only cells containing exactly one variable heavy (V_H_) and variable light (V_L_) chain were considered. The germline node represents the unmutated reference sequence determined by 10X Genomics cellranger.

**Figure S11.**
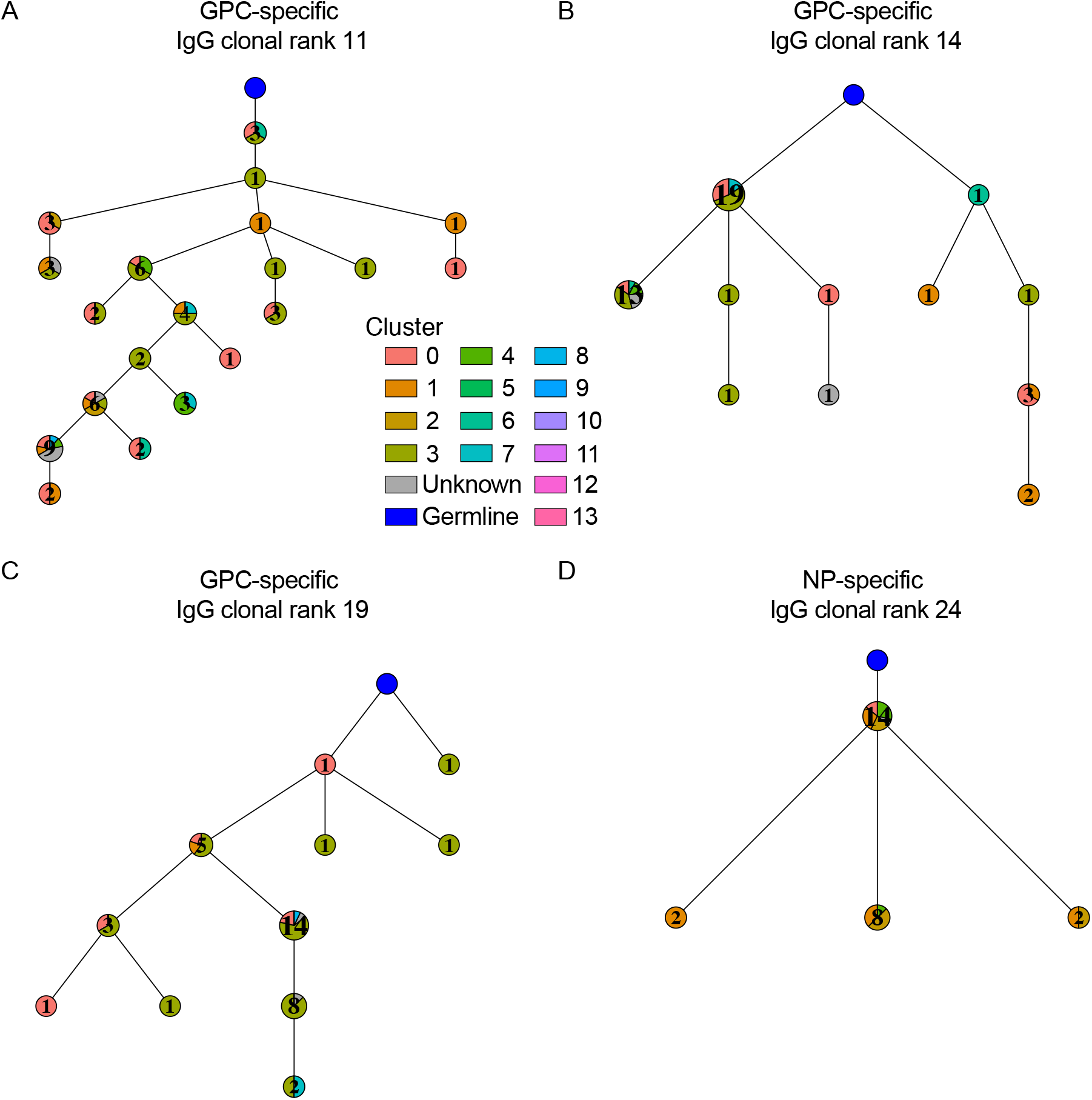
Virus-specific mutational networks displaying transcriptional cluster distribution. Nodes represent unique antibody variants (combined V_H_+V_L_ nucleotide sequence) and edges demonstrate sequences with the smallest separation calculated by edit distance. Node color corresponds to transcriptional cluster distribution for each cell. The size and label of the nodes indicate how many cells express each full-length antibody variant. Clone was determined by grouping those B cells containing identical CDRH3+CDRL3 amino acid sequences. Only cells containing exactly one variable heavy (V_H_) and variable light (V_L_) chain were considered. The germline node represents the unmutated reference sequence determined by 10X Genomics cellranger.

**Figure S12.**
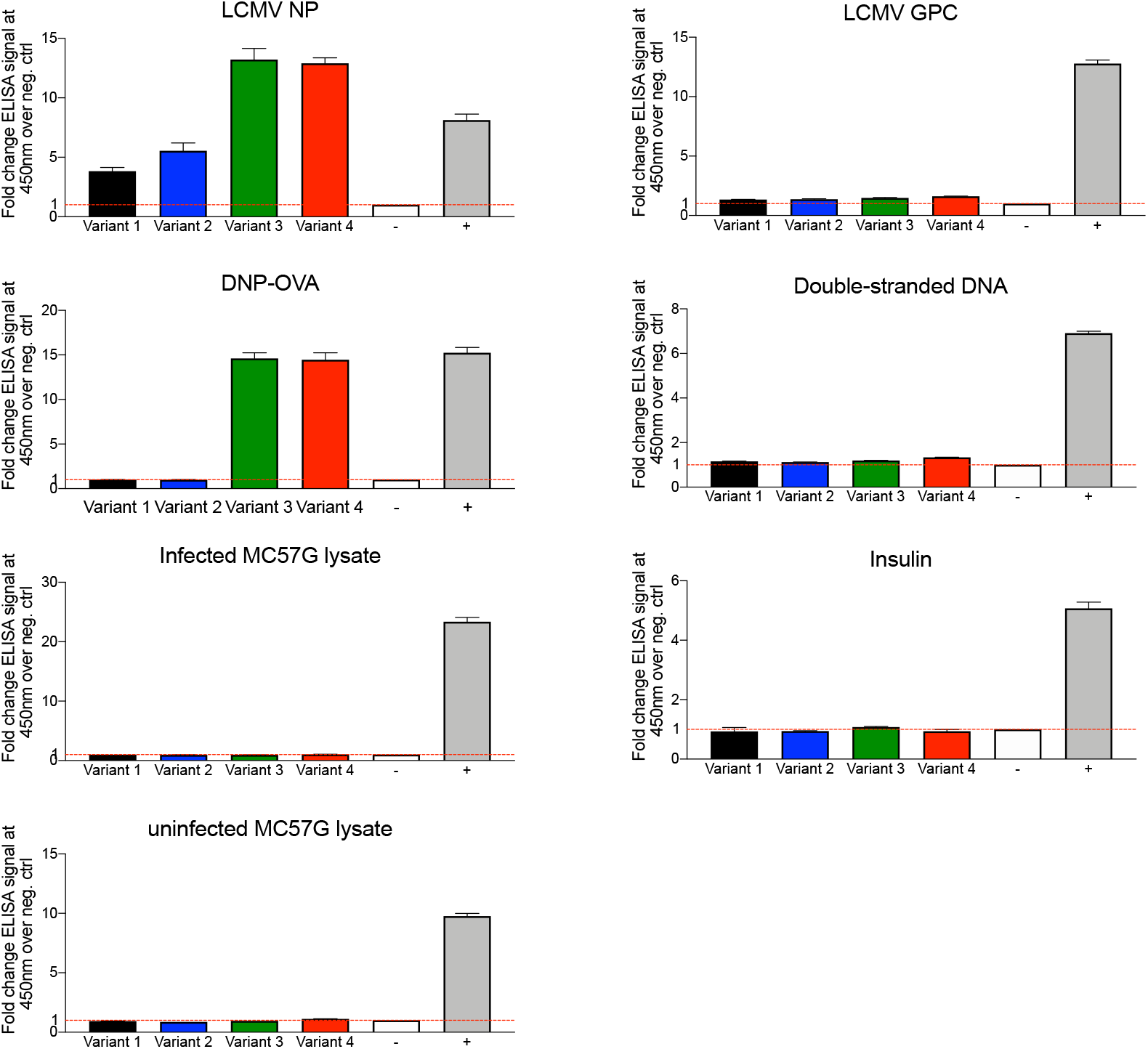
DNP-OVA cross-reactive antibody variants do not react with other antigens and cell lysates. The ELISA signal of duplicate ELISA measurements at 450 nm is shown relative to a negative background control (red dotted line indicates background level). Control antibodies used are listed in the methods section.

